# Multidimensional proteomics identifies molecular trajectories of cellular aging and rejuvenation

**DOI:** 10.1101/2023.03.09.531951

**Authors:** Mario Leutert, Joe Armstrong, Anja R. Ollodart, Kyle Hess, Michael Muir, Ricard A. Rodriguez-Mias, Matt Kaeberlein, Maitreya Dunham, Judit Villén

**Author notes:** Correspondence (M.L.), (J.V.).

## Abstract

The declining capacity of cells to maintain a functional proteome is a major driver of cellular dysfunction and decreased fitness in aging. Here we assess the impact of aging on multiple proteome dimensions, which are reflective of function, across the replicative lifespan of *Saccharomyces cerevisiae*. We quantified protein abundance, protein turnover, protein thermal stability, and protein phosphorylation in mother yeast cells and their derived progeny at different ages. We find progressive and cumulative proteomic alterations that are reflective of dysregulation of complex assemblies, mitochondrial remodeling, post-translational activation of the AMPK/Snf1 energy sensor in mother cells, and an overall shift from biosynthetic to energy-metabolic processes. Our multidimensional proteomic study systematically corroborates previous findings of asymmetric segregation and daughter cell rejuvenation, and extends these concepts to protein complexes, protein phosphorylation, and activation of signaling pathways. Lastly, profiling age-dependent proteome changes in a caloric restriction model of yeast provided mechanistic insights into longevity, revealing minimal remodeling of energy-metabolic pathways, improved mitochondrial maintenance, ameliorated protein biogenesis, and decreased stress responses. Taken together, our study provides thousands of age-dependent molecular events that can be used to gain a holistic understanding of mechanisms of aging.

## Introduction

The age-dependent decline in the capacity of cells to maintain a functional proteome is a major driver of cellular dysfunction and results in increased frailty, decreased fitness, and rise in mortality at the cellular and organismal level^1–6^. Aging leads to extensive changes in gene expression, translation, homeostasis, metabolism, and signaling, all of which directly impact protein properties such as protein abundance, protein turnover, protein stability, and protein post-translational modifications ^4,5^. Protein abundance determines the composition of the proteome and its age-dependent changes cause alteration in cellular processes and pathways^7,8^. Protein turnover, the balance between protein synthesis and degradation, sets protein abundance and maintains proteostasis. Dysfunction of protein turnover leads to the accumulation of damaged proteins and is associated with several age-related diseases^9,10^. Protein stability is a function of protein structure, folding state, and interactions. Upon aging, protein stability is affected because proteins are exposed to changing environments and damage, undergo structural changes, and the machinery that maintains proteins soluble is changing^11–14^. Moreover, different post-translational modifications are widely affected by age^7,15,16^. Signaling systems that depend on protein phosphorylation, such as the target of rapamycin (TOR) and the AMP-activated protein kinase (AMPK) pathways, have central and conserved roles in aging and are able to extend lifespan when modulated^17,18^. Together, these highly interconnected protein properties define the functionalities of the proteome, and their progressive and cumulative dysregulation in aging leads to deterioration of proteome integrity^5,19^. With the advent of novel technologies that combine quantitative mass spectrometry with cell biological and biochemical methods, the systematic analysis of protein abundance, protein turnover, protein stability, and phosphorylation states is now possible^19–26^. Integrated analysis of multiple proteomic dimensions that are highly relevant in aging promises a more comprehensive understanding of deteriorating functions and homeostatic failure that would not be possible through the study of individual genomic, proteomic, or phenotypic features.

The budding yeast *Saccharomyces cerevisiae* has served as an important model organism to study the basic biology of aging and has been crucial in understanding conserved mechanisms involved in longevity such as TOR and AMPK (yeast homolog: Snf1) signaling and caloric restriction^27^. The yeast replicative lifespan (RLS) model is defined by the number of daughter cells that an asymmetrically dividing mother cell generates prior to senescence, with wild type yeast strains averaging about 20-25 generations^28,29^. Based on the asymmetric cell division, yeast mother cells retain and accumulate aging factors with each generation that lead to prominent cellular phenotypes and are thought to determine mother lifespan while rejuvenating daughter cells^30,31^. Aging factors include passively or actively segregated proteins, damaged or aggregated proteins, specific cellular structures, and extrachromosomal rDNA circles^32–39^. Old mother cells experience deficits in multiple dimensions of physiological performance, including loss of pheromone sensitivity, slower division cycles, loss of genome maintenance capacity, loss of sporulation ability, loss of asymmetric division capacity, defects in proteostasis, defects in pH homeostasis, mitochondrial dysfunction, and failure in their ability to sense glucose properly^40–47^.

Aging research is increasingly transitioning to studying how molecular mechanisms integrate at the systems level in order to understand complex concepts involved in aging like homeostasis, signaling networks, and resilience. Comprehensive studies at the genetic, metabolic, and phenotypic level allowed dissecting the aging process in yeast in great detail; however, it is barely understood how relevant proteome dimensions are impacted by age^48–53^. Studying replicative aging in yeast requires the physical separation of mother cells from their progeny. Manual manipulation^54^ or microfluidic traps^52^ are typically used to quantify the median lifespan of a population of hundreds to thousands of individual mothers. Although these approaches are well suited for the quantification of RLS and have revealed a wide variety of aging phenotypes^48,53^, they are limited by their inherently laborious nature and cannot produce the sizable quantities of aging cells required for proteomics. An alternative approach to isolate aging mother cells is to label the cell walls of a population using magnetic beads^41,50^. During budding, daughter cells have newly formed cell walls, and beads remain with the mother cells, allowing them to be separated. Hendrickson et al.^49^ furthered this approach by incorporating magnetically labeled starting cells into a chemostat, which maintains a constant environment and retains the starting population by an external magnet.

Here, we combined aging of yeast under highly controlled conditions in chemostats with cutting-edge multidimensional proteomics technologies to create a quantitative atlas of protein abundance, turnover, thermal stability, and phosphosites across RLS in mother and daughter cells, that can be explored in our web-based analysis tool (https://rlsproteomics.gs.washington.edu). We integrated the multidimensional proteomic data in a single model and dissected gradual age-dependent changes to organelle morphology and functions, mapped trajectories for various protein complexes, and profiled the activation states of kinase-signaling pathways. We uncovered pervasive proteomic alterations that we could connect to known aging hallmarks, asymmetric protein segregation, and a shift from biosynthetic to energy-metabolic processes at multiple levels. Our findings further extend the asymmetric segregation concept between mother and daughter cells to protein complex assemblies, protein phosphorylation, and signaling pathway activities. We mechanistically dissect the age-dependent remodeling of the mitochondrial proteome and the underlying energy-metabolic signaling. We identify asymmetric activation of the AMPK/Snf1 as a key event, leading to glucose derepression in mother cells. We follow up on the observed energy-metabolic pathway remodeling in aging by analyzing age-dependent proteomic changes in a genetic *HXK2* deletion (*hxk2Δ*) caloric restriction model. We observe reprogrammed energy-metabolic pathways in young *hxk2Δ* cells, which results in limited proteomic remodeling of the same pathways in aging, possibly conserving resources during aging. Furthermore, caloric restriction enhanced the maintenance of proteins involved in biogenesis, reduced morphological changes to the mitochondria, and reduced induction of stress responses.

We provide a systems-level view of how the yeast proteome is affected by aging and anticipate that our resource will be useful to scientists from diverse fields and accelerate research into the basic biological mechanisms of aging, which could lead to improved clinical interventions for age-related diseases and conditions.

## Results

### A framework to derive replicative aged yeast mother and daughter cells for proteomics

To adapt the replicative aging yeast model for quantitative proteomics, we had to overcome the following challenges: i) separate mother cells from daughter cells; ii) obtain mother cells with a defined and narrow age range; iii) ensure the aging process is not influenced by sample handling or changing environments; and iv) collect enough old mother cells to accommodate the sensitivity of various proteomic assays. We used the concept of labeling yeast cell walls with functionalized magnetic beads to separate mother and daughter cells^41^. Previous magnetic bead labeling approaches for yeast cells relied on biotinylation of cells followed by labeling with magnetic streptavidin functionalized beads^41,49,50^. This approach includes outgrowth in batch culture and inefficient recovery of labeled cells, is prohibitively expensive for large amounts of cells, and requires the addition of biotin precursors to prevent biotin starvation^49^. To address these issues, we devised a novel labeling method that involves directly crosslinking carboxylate-modified magnetic beads to free amines on the surface of yeast cells in a single step (Extended Data Fig 1a). Our approach uses mild conditions for crosslinking (30 min in phosphate buffered saline at room temperature), does not require outgrowth, is cost-effective, and results in uniformly labeled yeast with 1-4 beads per cell on average (Extended Data Fig 1b). To determine the effects of beading on RLS, we performed microdissection experiments^48^ on beaded cells, mock beaded cells, and cells that did not undergo the beading procedure. We found that there was no difference in RLS between beaded and mock beaded cells (Extended Data Fig 1c,d). We also found that around 20% of the beaded and mock beaded population does not appear to divide when transferred to solid media (Extended Data Fig 1c,d).

**Extended Data Figure 1.**
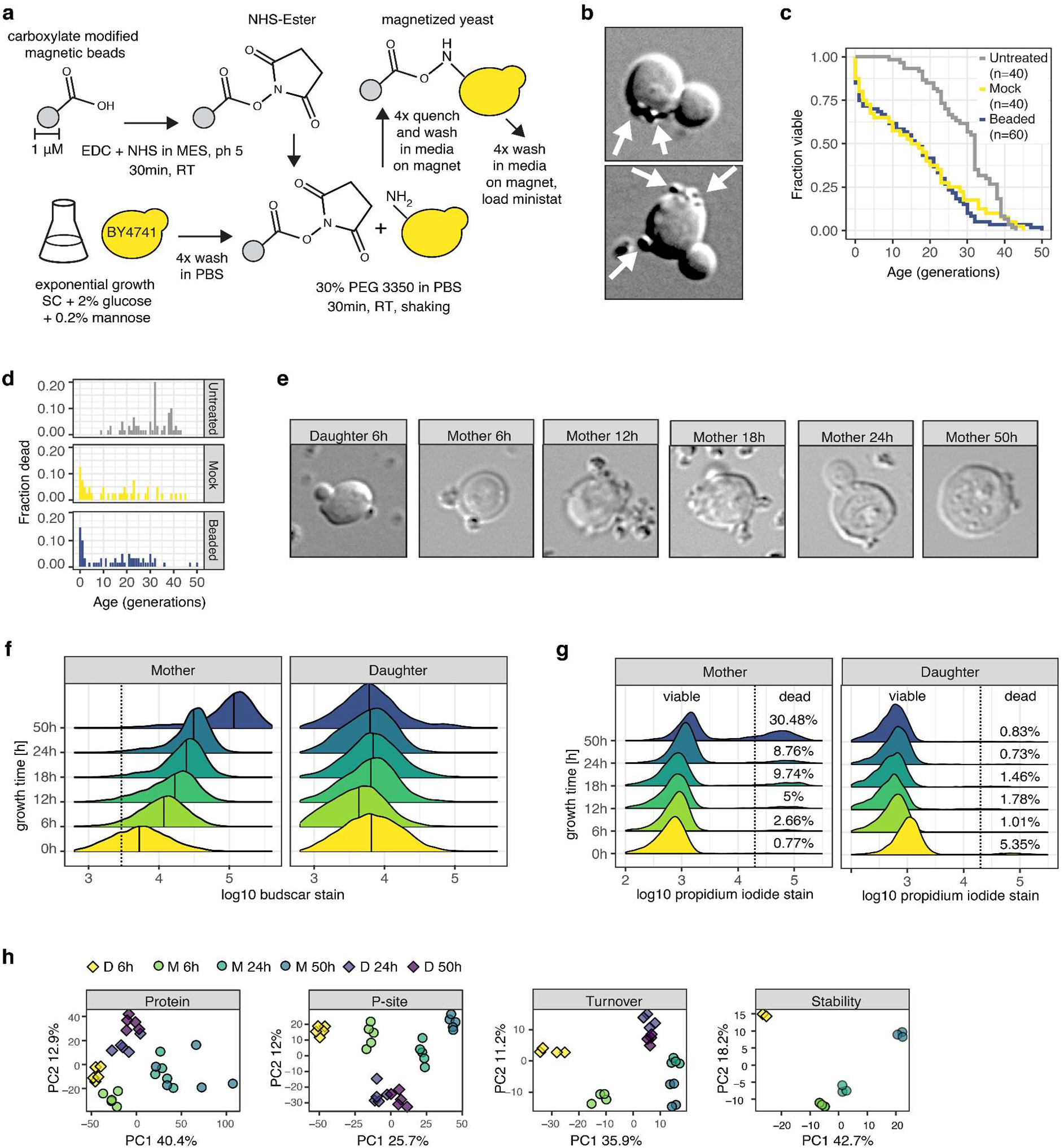
Magnetic beading of yeast cells. **A)** Overview of protocol to activate magnetic beads, prepare yeast cells and cross-link beads to yeast cell surface. **B)** Representative microscopic brightfield picture of two freshly beaded cells with two and three crosslinked beads respectively (indicated with white arrow). **C)** RLS survival curves for untreated, beaded and mock beaded cells determined by microdissection assay. **D)** Histogram of death occurrence across age for same cells as in F). **E)** Microscopic brightfield pictures of cells obtained from the MADs at different timepoints shown in Fig. 1B. **F)** Density plots of flow cytometry quantified WGA bud scar stain fluorescence of cells obtained from the MADs at different timepoints. Same samples as in Fig 1c. **G)** Density plots of flow cytometry quantified propidium iodide cell death stain fluorescence of same samples as in D). **H)** Principal component analysis for all proteomic dimensions across all samples.

To efficiently obtain mother cells of defined ages that were grown in a constant environment, we implemented the miniature-chemostat aging devices (MADs) developed by Hendrickson et al.^49^. Beaded yeast cells are directly loaded into a miniature-chemostat housed in a strong neodymium magnet. Fresh medium and filtered air are delivered at high flow rates via a peristaltic pump, and daughters are washed away (Fig 1a). Thus, levels of nutrients, aeration, and pH are constant throughout the experiment. To profile mother and daughter cells across RLS, entire MADs are harvested at different timepoints, mothers are separated from their non labeled direct daughter cells and both populations are collected. We used bud scar staining and manual bud scar counting by microscopy, as well as fluorescence quantification by flow cytometry, to confirm constant aging and high purity of yeast mothers and daughters collected from the MAD at multiple timepoints (Fig 1b-d, Extended Data Fig 1e,f). Yeast mothers grown in the MAD for 50 hours have a high average age of 18.2 bud scars and a 40% increase in doubling time (Fig 1e). Viability of mother cells grown for 24 hours in MADs was 90% and decreased to 70% after 50 hours (Extended Data Fig 1g).

**Figure 1.**
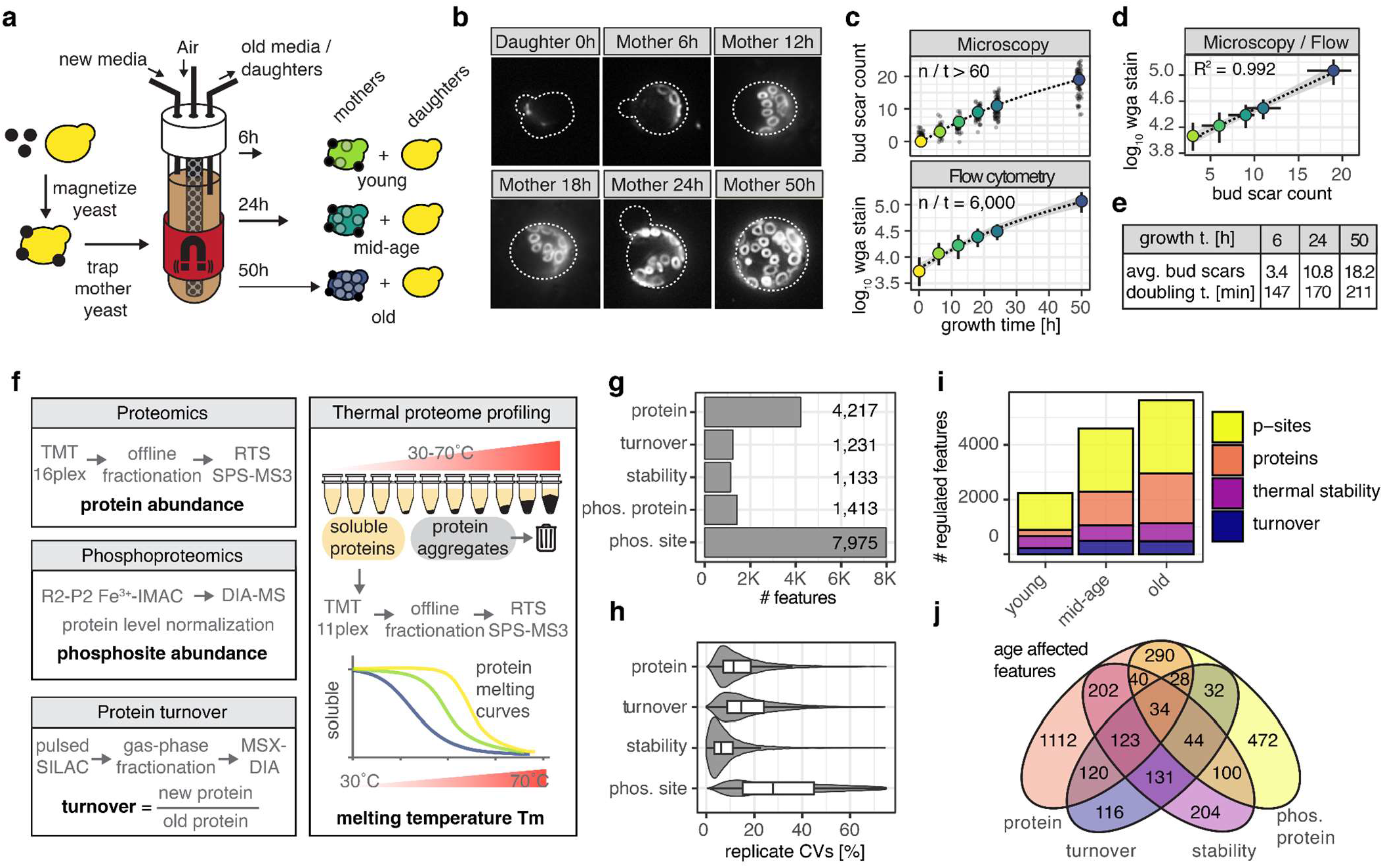
Multidimensional proteomics of replicative aged yeast. **A)** Approach to obtain replicative aged yeast cells using miniature-chemostat aging devices (MADs). **B)** Representative microscopic pictures of WGA stained bud scars on cells obtained from the MADs at different timepoints. **C)** Age determination by WGA bud scar staining and manual counting using microscopy or by WGA fluorescence quantification using flow cytometry. **D)** Correlation of bud scar quantifications between flow cytometry and manual counting. **E)** Determined average bud scar count and doubling time for 3 harvesting timepoints. **F)** Experiments to determine multiple protein properties proteome-wide. TMT = tandem mass tag, RTS = real time search, SPS = synchronous precursor selection, R2-P2 = Rapid robotic-phosphoproteomics, DIA = data independent acquisition. **G)** Count of quantified and reproducibly measured protein properties across all samples. **H)** Coefficient of variation (CV) for different measurements. **I)** Count of significantly regulated protein properties across different mother ages versus young daughters. **J)** Venn diagram of proteins that show significant regulation in abundance, phosphorylation, stability and turnover across all mother ages versus young daughters.

Taken together, we established a framework that allows for the separate collection of large amounts of yeast mother cells and their daughter cells at high purity across yeast RLS up to old age. Based on our findings we choose to harvest MADs and collect mother and daughter cells for proteomic assays at 6h for young cell populations, at 24h for middle-aged populations, and at 50h for old populations.

### A proteomic atlas of protein abundance, turnover, thermal stability, and phosphorylation across RLS

We collected and characterized yeast mother cells and their daughter cells from three age groups: 6 hours of growth for young cells (3.4 average bud scars), 24 hours for middle-aged cells (10.8 average bud scars), and 50 hours for old cells (18.2 average bud scars). Daughter cells from young mother cells (6 hour collection) were set as the baseline proteome for young cells. Daughter cells that were collected from across RLS (6, 24, and 50 hour collections) were then used to study asymmetric division and rejuvenation potential with age. We acquired proteome-wide measurements of multiple protein properties across the different age groups and cell populations using the following mass spectrometry (MS) based assays (Fig 1f): i) To identify changes to proteome composition, we used multiplexed proteomics and quantified the abundance of ∼4200 proteins (Supplementary Table 1); ii) To gain insights into changes of protein synthesis, degradation, and asymmetric partitioning we performed pulsed metabolic labeling with heavy isotope lysine in the MADs and determined turnover rates for ∼1200 proteins^24,55^ (Supplementary Table 2); iii) To profile protein transitions reflective of folding-states, environmental conditions, substrate binding, protein-protein interactions, and spatial rearrangements^21^, we performed thermal proteome profiling (TPP)^20^ and determined thermal stability values for ∼1300 proteins (Supplementary Table 3); iv) To map activated kinases and regulated signaling pathways, we performed phosphoproteomics^25,56^ and quantified ∼8000 phosphosites for ∼1400 phosphoproteins across all samples (Supplementary Table 4). Overall, we achieved deep, precise, and reproducible proteome-wide measurements across all dimensions and conditions (Fig 1g,h). Principal component analysis for each proteomic dimension revealed separation by principal component one based on age and principal component two based primarily on differences between mother and daughter cells (Extended Data Fig 1g). We used linear modeling to identify protein properties that change in old mother cells versus young daughter cells, and we discovered a steady increase in changing properties with age, with over 5000 protein properties differently regulated in old mother cells (Fig 1i, Supplementary Table 5). The majority of the changes occurred from young to middle-aged mother cells. We discovered that different proteomic measurements often captured regulation of protein properties that were regulated independent of properties of the same protein, underscoring the benefit of acquiring multiple proteomic dimensions at the same time (Fig 1j).

Taken together, we have developed a comprehensive and high quality proteomic atlas of protein abundance, turnover, stability, and phosphosites in yeast mother cells and their direct daughter cells at different ages. The atlas can be interactively explored through our web-based tool (https://rlsproteomics.gs.washington.edu) and interrogated to identify age-dependent or mother-daughter specific changes across structural, biochemical, cellular, and signaling dimensions.

### Pervasive proteomic alterations are related to aging hallmarks

Using our multidimensional proteome atlas, we first sought to correlate proteome changes with recognized markers of yeast RLS, specifically focusing on changes related to asymmetric cellular partitioning, vacuole size, pH homeostasis, loss of mitochondrial membrane potential, and protein damage (Fig 2a)^39,57^.

**Figure 2.**
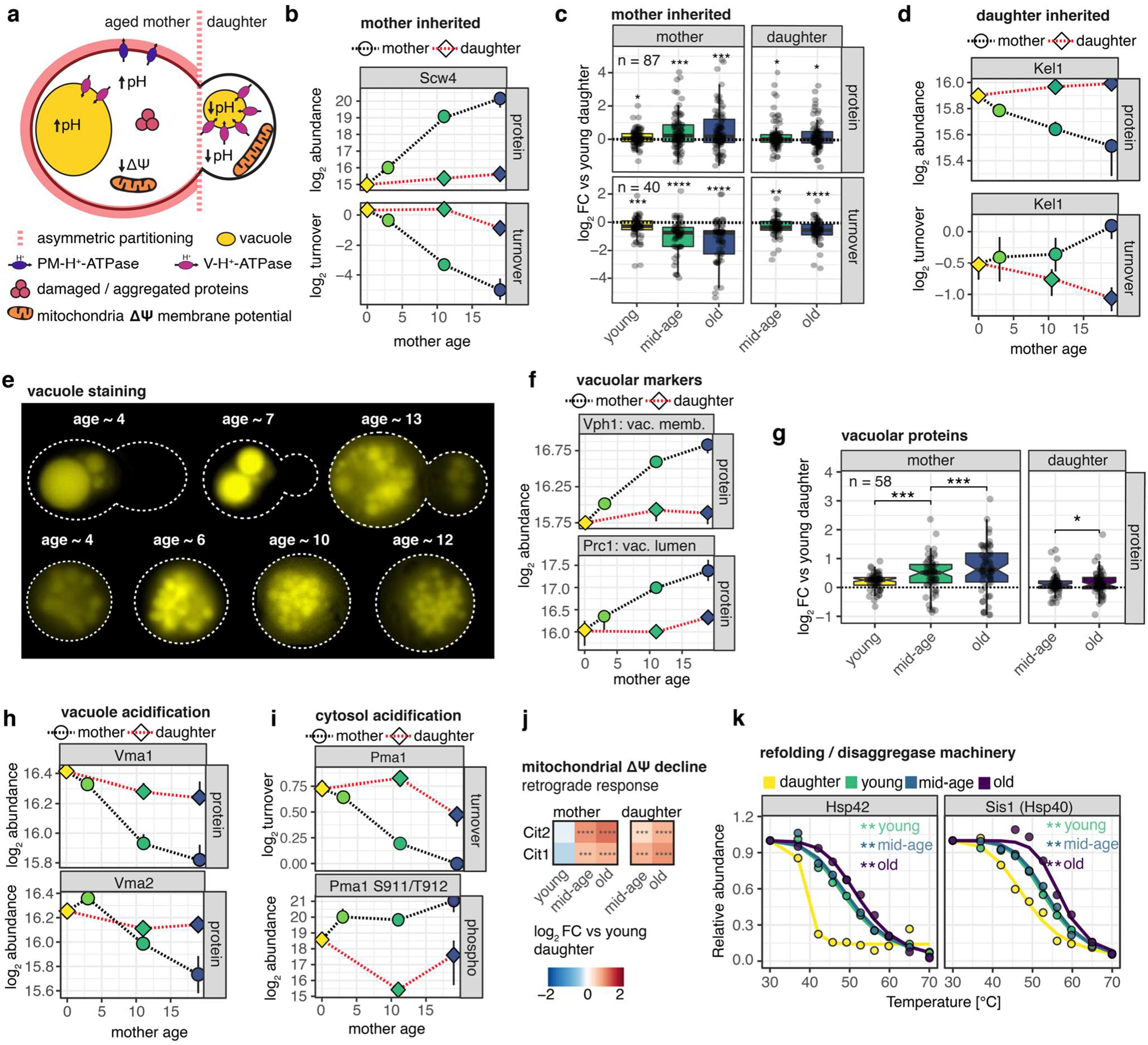
Pervasive proteomic signatures of aging hallmarks in mother cells. **A)** Reported aging hallmarks expected to affect the proteome. **B)** Protein abundance and turnover for the cell wall protein Scw4, which has been reported to be asymmetrically inherited by mother cells. **C)** Log_2_ fold changes of protein abundance and turnover for proteins previously reported to show asymmetric segregation to mother cells. One-sided t-tests were performed to test whether group mean is different from 0 and statistical significance is indicated by: * p< 0.05, ** p < 0.01., *** p < 0.001, **** <p 0.0001. **D)** Protein abundance and turnover for Kel1, which has been reported to be asymmetrically segregated to daughter cells. **E)** Vacuolar staining of mother cells at different ages derived from MADs. **F)** Protein abundance for Vph1, a vacuolar membrane marker, and Prc1, a vacuolar lumen marker. **G)** Log_2_ fold changes of protein abundance for vacuolar proteins. Group tests were performed using a nonparametric Wilcoxon test. Statistical significance is indicated by: * p< 0.05, *** p < 0.001. **H)** Protein abundance for Vma1 and Vma2 subunits of the V-ATPase. **I)** Turnover level of PM-ATPase Pma1 and abundance of activating phosphosites S911 and T912. **J)** Protein abundance log_2_ fold changes for proteins induced by the retrograde response. **H)** Melting curves for proteins involved in disaggregating unfolded proteins. Statistical significance versus young daughter cells is indicated (** q-value < 0.01).

Proteins that are asymmetrically retained by mother cells gradually accumulate after each division^31^. Several of these proteins have previously been identified using microscopy, flow cytometry, and proteomics^33–36^. We expected that mother-retained proteins would result in pronounced age-dependent and mother-specific increases in protein abundance and decreases in turnover. This trend was validated by plotting abundance and turnover of the well-known mother cell inherited cell wall protein Scw4 (Fig 2b). Next, we considered proteins that had at least two instances of mother cell inheritance in prior studies and found good agreement with the hypothesized protein abundance and turnover profile in our data (Fig 2c). Remarkably, we also discovered that in daughter cells from old mother cells, asymmetric partitioning gradually begins to break down for a small number of proteins (Fig 2c). Although asymmetrically inherited proteins by daughter cells have been reported too^33^, they are difficult to identify because they do not accumulate, and it is unknown whether they play a role in aging. We observe depletion and rapid turnover in mother cells for a few documented daughter cell-inherited proteins, mostly localizing to bud structures, but we do not detect a general pattern (Fig 2d, Extended Data Fig 2a).

**Extended Data Figure 2.**
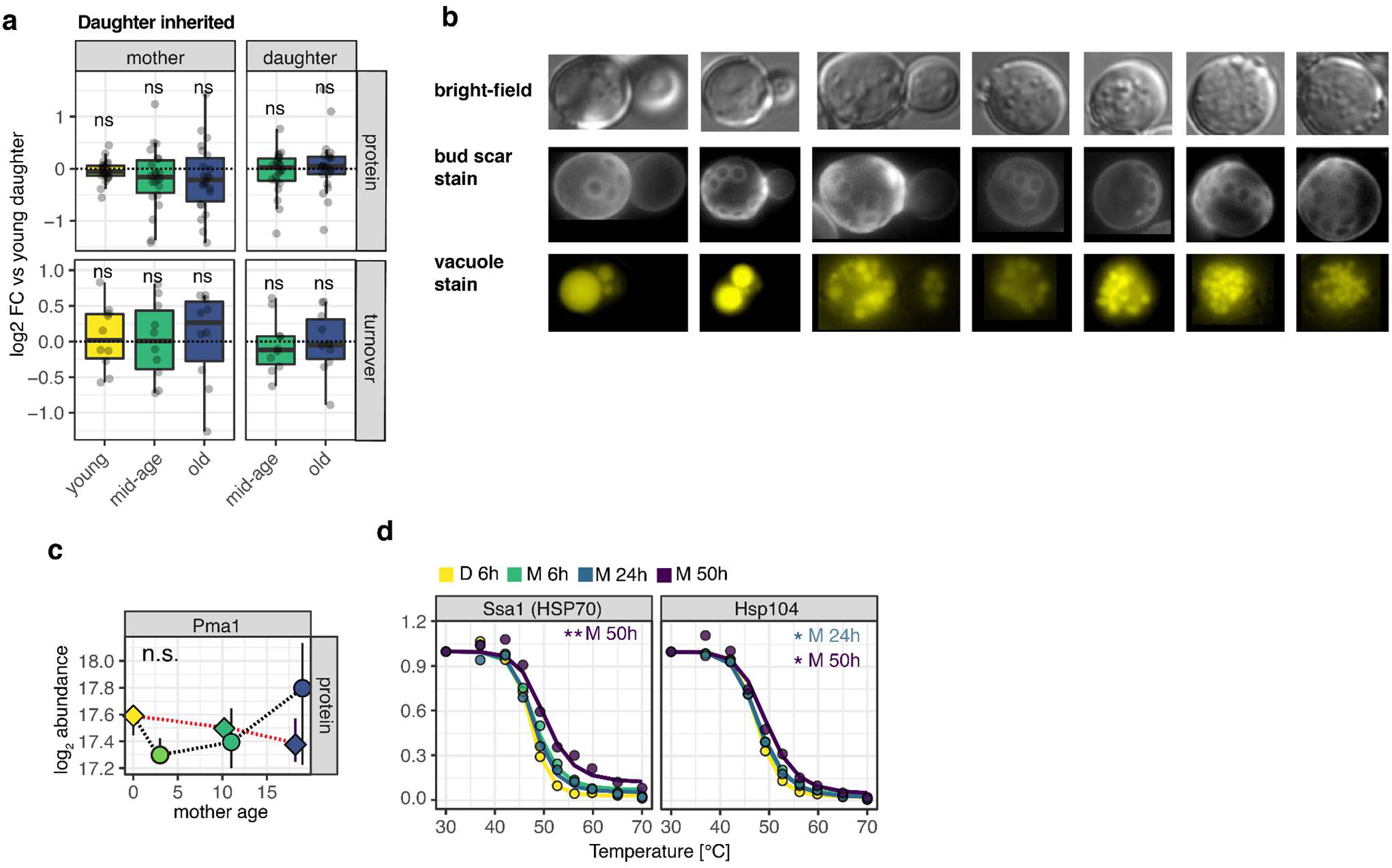
Signatures of aging hallmarks. **A)** Fold changes of protein abundance and turnover for proteins previously reported to show asymmetric segregation to daughters cells^33^. One-sided t-tests were performed to test whether group mean is different from 0, ns = non significant. **B)** Bright field, bud scar and vacuole staining of mother cells shown in Fig 2e. **C)** Protein abundance of Pma1. **D)** Melting curves for proteins involved in disaggregating unfolded proteins. Statistical significance versus young daughter cells is indicated (* q < 0.05, ** q < 0.01).

Increase in vacuole size in relation to total cell size is a prominent characteristic of aged yeast cells that is partially caused by asymmetric partitioning at the organelle level^58^. Vacuolar morphology was reported to be heterogeneous from cell to cell and across lifespan with both fused and fragmented vacuoles occurring late in old cells^59^. By using microcopy, we qualitatively validated increased vacuolar loads in old mother cells derived from MADs (Fig 2e, Extended Data Fig 2b). We were able to corroborate this increased vacuolar load using our proteomic data, where we observed an age-dependent, mother-specific quantitative increase of widely utilized vacuolar membrane and lumen protein markers (Fig 2f). Analysis of the abundance of 58 vacuole specific proteins further confirmed a mother-specific, age-dependent increase in vacuolar proteome abundance compared to the rest of the proteome (Fig 2g)^60^.

Another hallmark of yeast aging and RLS is the loss of vacuolar and cytosolic acidity in mother cells, which are subsequently re-acidified in daughter cells^45,61^. Essential for maintaining vacuolar acidity is Vma1, the catalytic subunit of the vacuolar-type ATPase (V-ATPase). While *VMA1* gene deletion causes premature maternal cell death, *VMA1* overexpression lengthens RLS^45,62^. Consistent with these phenotypes, we find age-dependent decline of Vma1 and Vma2 protein levels in mother cells, but not daughter cells (Fig 2h). Cytosolic pH is mainly controlled by the H^+^-ATPase Pma1. However, unlike Vma1 and Vma2, we find that the cytosolic pH asymmetry cannot be explained by changes in overall Pma1 protein levels (Extended Data Fig 2c). Instead, we find that Pma1 has a slower turnover in mother cells when compared to daughter cells, and, more strikingly, activating tandem phosphosites Ser-911 and Thr-912 on Pma1^63^ are strongly upregulated in old mother cells, while they largely remain dephosphorylated in daughter cells (Fig 2i). This indicates that asymmetric cytosolic pH regulation by Pma1 may be post-translationally regulated and illuminates a potential novel mechanism of asymmetric phosphorylation.

Mitochondrial dysfunction due to declining membrane potential was reported in aging yeast^45,64^. Mitochondrial dysfunction triggers retrograde signaling which induces a nuclear gene expression program that aims to reconfigure metabolism to account for mitochondrial defects ^65^. We find that both Cit1 and Cit2 —prototypical targets of the retrograde response^65^— are significantly upregulated in middle-aged and old mother cells as well as in their respective daughter cells (Fig 2j). This shows that retrograde signaling is activated early in the aging process and is passed on to daughter cells.

Damaged, misfolded, and aggregated proteins are known aging factors^37,66^. It has been demonstrated that aged yeast mother cells inherit protein aggregation deposits that are nucleated by Hsp42 oligomerization and partially resolved by the Hsp104/70/40 chaperone protein refolding machinery^66^. We identified a strong age-dependent increase in Hsp42 thermal stability, which is suggestive of its oligomerization (Fig 2k). Sis1 (a Type II HSP40 co-chaperone) also exhibits a strong age-dependent increase in thermal stability, which may be attributed to its propensity to bind misfolded proteins, and hence stabilization upon increased substrate binding in old cells with more misfolded proteins (Fig 2k). Hsp104 and Hsp70 are also stabilized in the old mother cells (Extended Data Fig 2d). Interestingly, the stability of refolding chaperones increased in young mother cells compared to their daughter cells, was similar in young and middle-aged mother cells, and increased again in old mother cells. Occurrence of broad structural changes already in young mothers is consistent with a recent structural proteomics study ^14^. These observations suggest that the primary differences in aggregate formation occur between mother and daughter cells (nucleation and asymmetric inheritance of aggregates) and later in life (possibly increased aggregation due to a decline in proteostasis).

Taken together, we discovered strong correlations between our multimodal datasets and established hallmarks of aging. We further characterize these aging processes by providing details on the dynamics of different protein properties, defining their time of onset, proteome-wide impact, and transmission to daughter cells.

### Age scores prioritize age-dependent protein changes across proteomic dimensions

We aimed to integrate the four proteomic data modalities into a single model to prioritize age-dependent changes of protein properties across proteomic dimensions. We achieved this by performing temporally-informed data dimensionality reduction across proteomic data modalities (for young daughters, young mothers, middle-aged mothers, and old mothers) using multi-omic factor analysis as implemented in the MEFISTO framework^67^. In total, this framework generated four factors, which captured the biological variability encoded across proteomic and aging dimensions. We found that factor 1 captured protein properties (abundance, turnover, stability, and phosphosites) that gradually vary as a function of age and explained the majority of the biological variance across all data modalities (Fig 3a, Extended Data Fig 3a). Factor 2 captured protein properties that differ between young mothers and their daughter cells; factor 3 identified alterations in middle-aged mothers; and factor 4 identified inherent variation in old mothers (Extended Data Fig 3b). The model provides mapping between the latent factors and all measured protein dimensions. To assess age-dependent changes uniformly, we retrieved factor 1 weights for all protein properties and termed them age scores (Supplementary Table 6). Age scores significantly improve our capacity to examine age-dependencies of protein properties across proteomic dimensions by scoring whether a protein property is increasing (positive age score) or decreasing (negative age score) as a function of age (Fig 3b). Age effects across proteomic dimensions are mostly orthogonal, with negative correlations for protein abundance versus turnover and phosphorylation (Extended Data Fig 3c). Numerous well-known mother-inherited proteins, such as the cell wall proteins Scw4 and Scw10 or the long-lived protein Hsp26, rank among the most positive age scores for abundance and most negative for turnover (Extended Data Fig 3d). Age scores are incorporated in our web-based analysis tool (https://rlsproteomics.gs.washington.edu).

**Figure 3.**
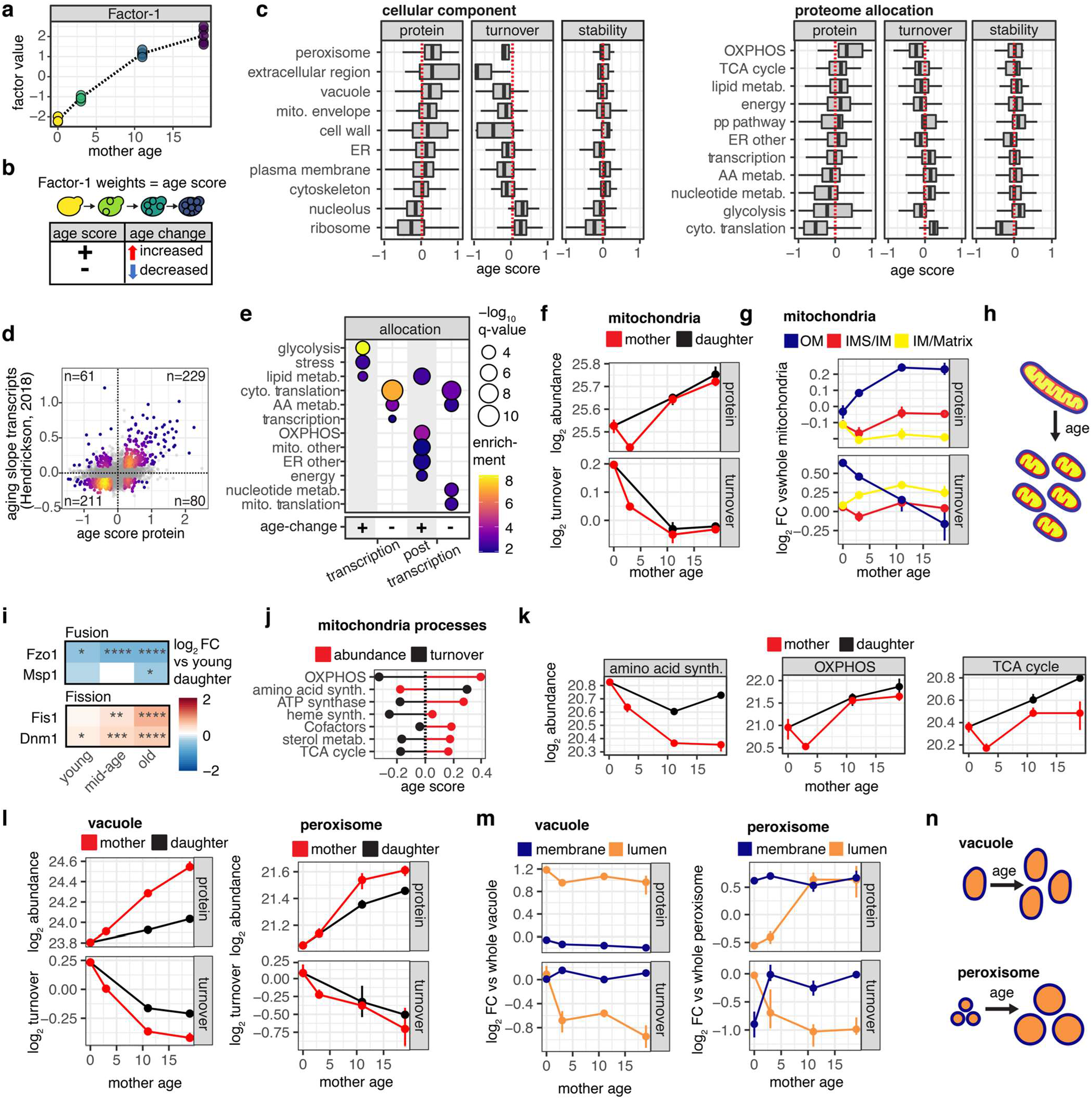
Impact of age on organelle function and morphology. **A)** Factor 1 from multi-omic factor analysis captured age dependent variation across samples. **B)** Factor 1 weights of protein properties across different dimensions are extracted and used as age scores to prioritize protein properties that are affected by age. **C)** Boxplot of age scores for abundance and turnover protein properties for cellular compartments and proteome allocation annotations. ER = endoplasmic reticulum, OXPHOS = oxidative phosphorylation, pp = pentose phosphate, AA = amino acid. **D)** Scatterplot of transcript age changes^49^ versus protein abundance age scores. Colored points represent transcripts and proteins that are regulated upon aging on the transcript and protein level, colored by density, and count in each quadrant is indicated. **E)** Significantly enriched proteome allocation terms for age dependent changes that occur at the transcriptional and/or at the post-transcriptional level. **F)** Summed abundance and average turnover of mitochondrial proteins in mother and daughter cells. **G)** Fold change of median abundance and turnover for mitochondrial proteins localized to outer membrane, the intermembrane space or the matrix versus the average of all mitochondrial proteins. **H)** Graphic representation of mitochondrial fragmentation that could explain observed changes to suborganelle mitochondrial proteomes. **I)** Log_2_ fold changes for key proteins mediating mitochondrial fusion and fission. **J)** Median abundance and turnover age scores for mitochondrial proteins functions. **K)** Median abundance and turnover for selected mitochondrial protein functions. **L)** Summed abundance and average turnover of vacuolar and peroxisomal proteins in mother and daughter cells. **M)** Fold change of median abundance and turnover for vacuolar or peroxisomal proteins either localized to the membrane or lumen versus the all annotated proteins of the respective organelle. **N)** Model of organelle morphological changes that could explain observed differential changes to spatial suborganelle proteomes.

**Extended Data Figure 3.**
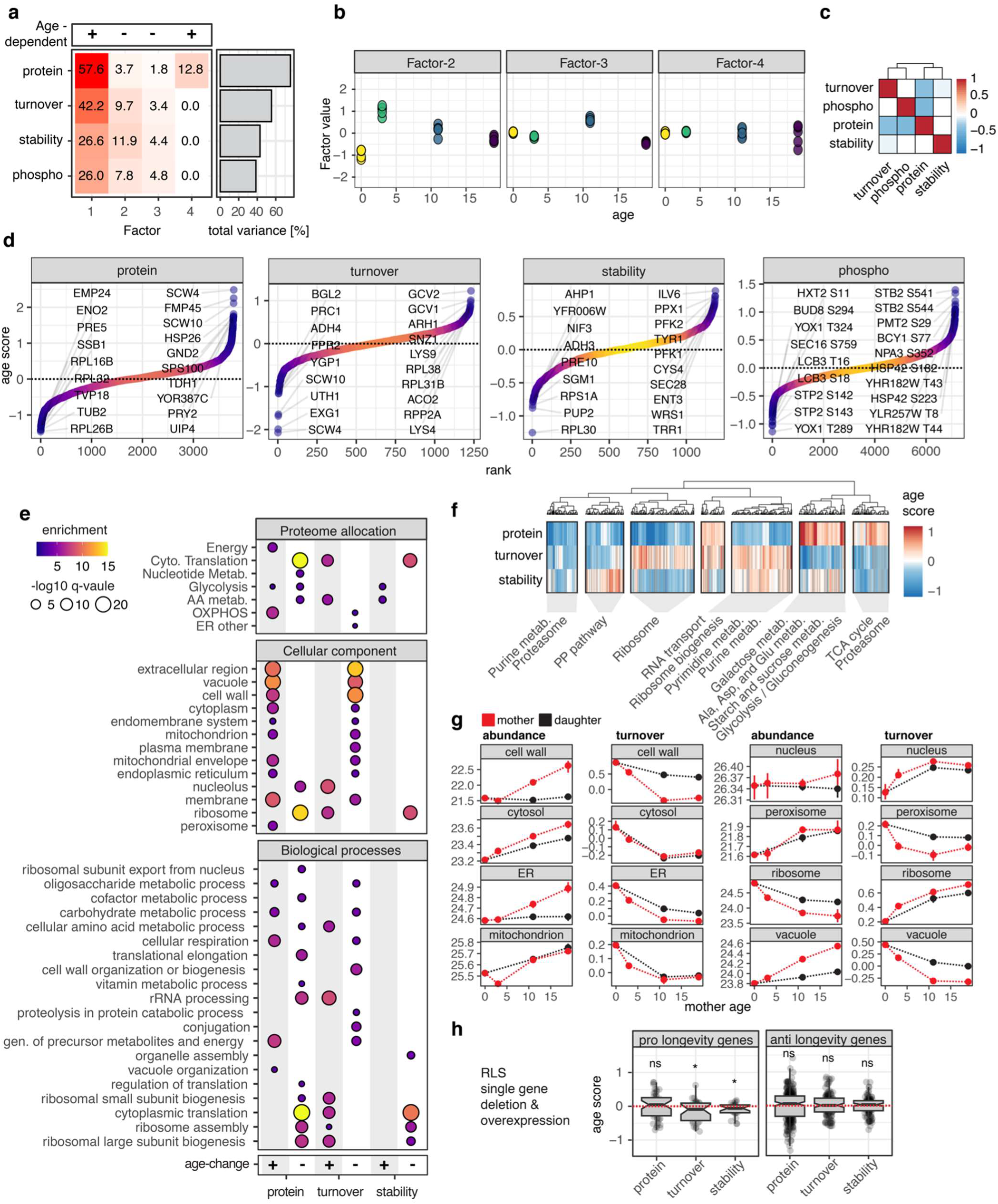
Age scores prioritize age-dependent protein changes. **A)** Unsupervised integration and dimensionality reduction by factor analysis across proteomic dimensions and age using MEFISTO^67^. Captured variance per factor and data modality is shown in heatmap and total captured variance of the four factors in the bar plot to the right. It is indicated if a factor identified temporal relationship (age) between samples. **B)** Factor values for Factor 2-4 versus age. **C)** Heatmap of person correlation of age scores for proteins across different proteomic data modalities. **D)** Rank plots of age scores for all data modalities. Protein features with highest and lowest age scores are indicated. **E)** Age score enrichment analysis for different gene annotations. **F)** Heatmap of age scores for proteins that have at least two measured dimensions and at least one age score with an absolute value > 0.2. Hierarchical clustering was performed, 7 main clusters were identified and enrichment analysis was performed within each cluster. Selected terms with a q-value <0.01 are indicated. **G)** Summed abundance and average turnover of proteins associated with different organelles in mother and daughter cells. **H)** Boxplot of age scores for all annotated pro- and anti longevity gene annotations derived from single gene deletion or overexpression. One-sided t-tests were performed to test whether group mean is different from 0 and statistical significance is indicated by: ns = non significant, * p< 0.05.

### Age-dependent remodeling from biosynthesis to energy metabolism

In order to comprehend how aging affects the proteome globally, we examined age score distributions and enrichments for protein abundance, turnover, and stability across subcellular localizations, functions, and pathways (Fig 3c, Extended Data Fig 3e-f). For proteins located in the cell wall, extracellular space, and vacuole, we identified strong patterns of increased abundance and decreased turnover that are consistent with an asymmetric mother inheritance mechanism. Protein subgroups related to energy metabolism and the peroxisome increased in abundance with age but were not significantly impacted by changes in turnover (Fig 3c, Extended Data Fig 3e-f). This type of regulation implies a general age-dependent increase in protein abundance that is similar in mother and daughter cells, consistent with increased gene expression. Proteins associated with cytoplasmic translation, amino acid metabolism, RNA polymerase, and nucleolus localization showed a significant decrease in abundance and increase in turnover. This pattern suggests these proteins are actively depleted and/or transcriptionally repressed as cells get older. Age-dependent protein stability changes revealed strong destabilization of proteins involved in ribosome biogenesis, ribosome assembly, translation machinery, and stabilization of proteins involved in amino acid and carbon metabolic processes.

Next, we aimed to determine the contribution of age-dependent transcriptional changes to the observed proteomic remodeling. We contrasted protein abundance age scores with age-dependent variations in transcript abundance determined by Hendrickson et al^49^. We found good agreement between age-dependent changes of proteins and transcripts, with 440 protein-transcript pairs changing in the same direction and 141 in opposite directions (Fig 3d). As suggested by the foregoing proteomic signatures, we could confirm transcriptional upregulation of energy metabolism and peroxisome genes as well as transcriptional repression of translation, amino acid metabolism, and RNA polymerase genes. Overall, regulation of proteins involved in glycolysis, stress responses, and transcription seemed to occur at the transcriptional level; regulation of proteins involved in mitochondrial processes and nucleotide metabolism at the post-transcriptional level; and regulation of cytoplasmic translation, amino acid, and lipid metabolism at both levels (Fig 3e). None of the aging hallmarks discussed were regulated at the transcriptional level, underlining the relevance of studying proteomic phenotypes of aging.

### Alterations of mitochondria, vacuole, and peroxisome morphology and function in aging

Our previous analysis showed that subcellular localization is a major factor that defines age-dependent changes to protein abundance and turnover. To obtain a systematic understanding of age-dependent subcellular proteome reorganization in mother and daughter cells, we compared the trajectories of total protein abundance and median turnover for various organelles and subcellular structures versus the whole cell (Extended Data Fig 3g). Different patterns revealed: organelle asymmetric segregation in mother cells, as well as age-dependent increases in mother cells for the vacuole, cell wall, and to a lesser extent the ER; age-dependent increases both in mother and daughter cells for the mitochondria, peroxisome, and cytosol; no change to the nucleus; and strong decreases in the ribosome in mother cells.

Mitochondrial dysfunction is a conserved aging process and the overall increase of mitochondrial proteins and reduced turnover in older cells was striking (Fig 3f). We investigated potential morphological restructuring of the mitochondria by comparing protein abundance and turnover changes of mitochondrial outer membrane (OM), intermembrane space (IMS), and matrix proteomes^68^ normalized to the whole mitochondrial proteome along age (Fig 3g). Interestingly, both the abundance and turnover data showed unequal age-dependent restructuring in different parts of the mitochondria (Fig 3g). While outer membrane proteins increased in abundance and decreased in turnover, the intermembrane space and matrix demonstrated the opposite trend, albeit to a lesser extent. This pattern suggested a morphological transition from larger (tubular) mitochondria to smaller (fragmented) mitochondria with larger surface, all in the light of an overall increase in mitochondrial load (Fig 3h). In further support of this model, we find age-dependent downregulation of Fzo1 and Msp1, which are required for mitochondrial fusion and upregulation of Fis1 and Dnm1, which are required for mitochondrial fission (Fig 3i). Fragmentation of mitochondria in old mother cells was observed before by microscopy ^45^. Mitochondrial functions were differentially affected by age: proteins associated with respiration and energy production increased, whereas proteins involved in amino acid and iron sulfur (Fe-S) cluster synthesis decreased (Fig 3j). Differential alterations in functional mitochondrial protein groups are indicative of adaptation to morphological changes and metabolic needs^69^. A decrease in connected matrix space would lead to impairment of mitochondrial membrane potential, and aged mother cells might compensate by increasing production of electron transport chain and proton pump proteins. Interestingly, we found asymmetric segregation of functional mitochondrial protein groups between mother and daughter cells. Proteins in the TCA cycle and proteins involved in amino acid metabolism were more abundant in daughter cells compared to mother cells (Fig 3k), suggesting the inheritance of fitter mitochondria by daughter cells. Mother-daughter cell asymmetry of specific mitochondrial functional protein groups might indicate asymmetric segregation of whole mitochondria based on its fitness.

Next, we investigated the morphological restructuring of the vacuole and peroxisomes, which are also strongly impacted by age (Fig 3l). We compared the abundance and turnover of organelle membrane proteins with lumen proteins normalized by all proteins annotated to the respective organelle (Fig 3m). Age had no effect on the vacuole’s abundance ratio of membrane or lumen proteins, while turnover of lumen proteins dropped quickly already in young mother cells (Fig 3m). In the peroxisome, the ratio of lumen protein abundance increased and turnover decreased (Fig 3m). These observations point to two different models of age-dependent organelle alterations: vacuole load increased overall, but the ratio of membrane to lumen remained constant, whereas peroxisomes grew in size with larger lumens (Fig 3n).

### Asymmetric protein segregation and its breakdown in old cells

Due to the importance of asymmetric division in yeast RLS, we sought to identify novel asymmetrically segregated proteins and determine the age-related limits of correct asymmetric segregation. Age scores indicating an age-dependent increase in protein abundance and a decrease in turnover correctly identified proteins previously reported to be inherited by mother cells (Fig 4a). In addition to 21 previously-known mother-inherited proteins, we discovered 198 new mother-inherited proteins with a threshold of absolute age scores >0.1 and complete measurements of abundance and turnover pairs (Supplementary Table 7). Of the mother-inherited proteins we identified, 30% belonged to the vacuole, cell wall, or plasma membrane, which have been reported to be asymmetrically distributed (Fig 4b). Interestingly, when we assess the segregation of these mother-inherited proteins in daughter cells derived from old mother cells, we found that 24% of proteins failed to undergo asymmetric segregation (Fig 4b). Asymmetric segregation breakdown primarily affected proteins in the cytosol and mitochondria, which might be attributed to inefficient diffusion barriers^39^ or changes in asymmetric segregation based on functional properties of mitochondria^70^ and might contribute to reduced rejuvenation of daughter cells from old mother cells^40^.

**Figure 4.**
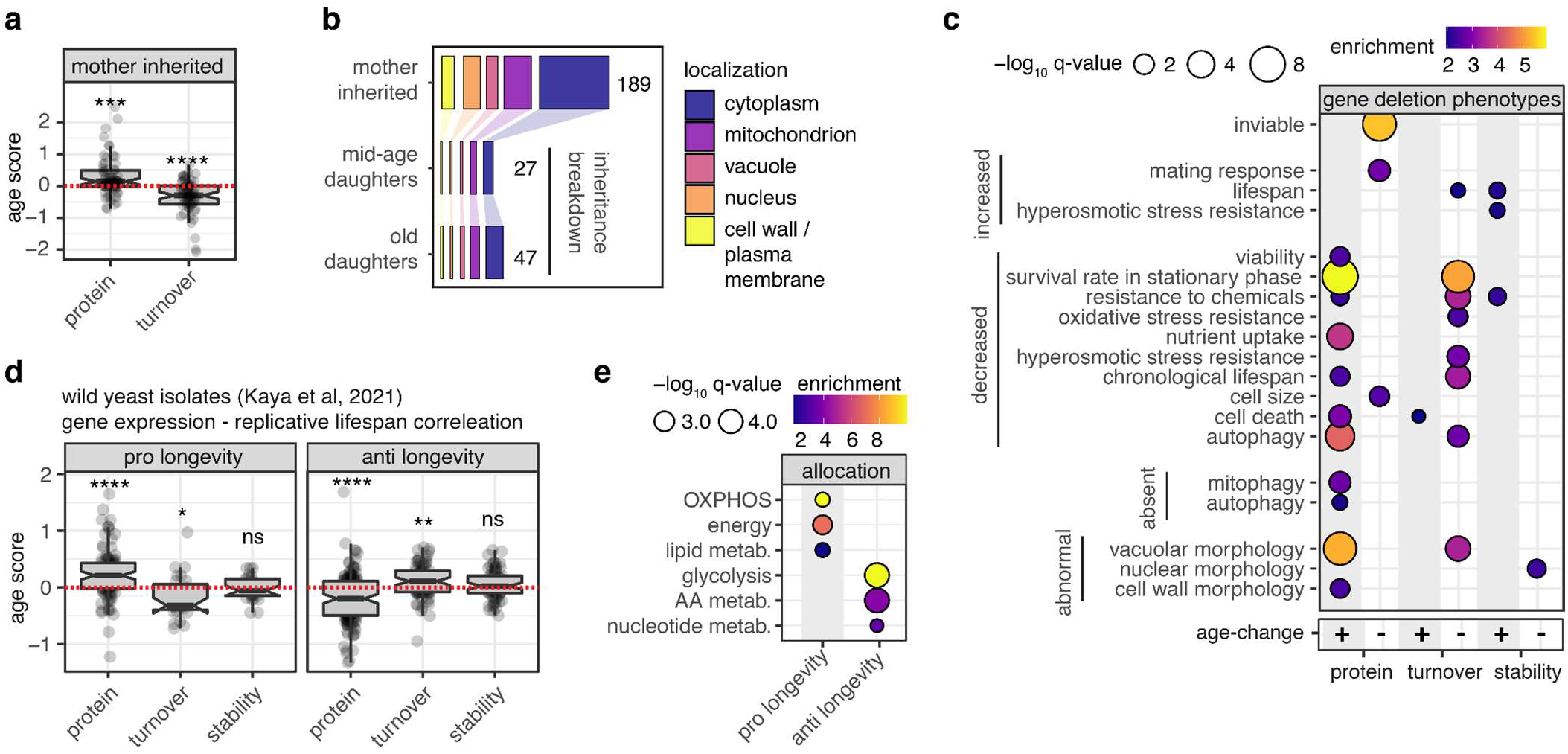
Asymmetric protein segregation, age-dependent phenotypes and longevity gene-expression associations. **A)** Protein abundance and turnover age scores for proteins previously reported to be inherited by mother cells. One-sided t-tests were performed to test whether group mean is different from 0 and statistical significance is indicated by: *** p < 0.001 and **** <p 0.0001. **B)** All identified mother cells inherited proteins annotated by subcellular localization. Proteins where asymmetric segregation breaks down in daughter cells from mid-aged and/or old mother cells. **C)** Enrichment for phenotype-gene deletion annotations in protein abundance age score. **D)** Boxplot of age scores for genes with positive or negative gene expression longevity correlation as determined by Kaya et al ^72^. One-sided t-tests were performed to test whether group mean is different from 0 and statistical significance is indicated by: ns =non significant, * p< 0.05, ** p < 0.01,**** <p 0.0001. **E)** Enrichment analysis of proteome allocation for proteins with positive age scores and positive gene expression longevity correlation and vice versa.

### Associations between age-dependent proteome changes and longevity

To better understand how age-dependent changes in protein properties influence cellular phenotypes, we calculated enrichments for protein abundance, turnover, and stability age scores across all yeast gene-deletion phenotype annotations (S288C background) (Fig 4c). Proteins with increased abundance and decreased turnover were enriched in genes that are involved in stationary phase survival, vacuolar morphology, autophagy, nutrient uptake, stress responses, and viability. Strikingly, proteins that decrease in abundance with age were enriched for essential genes (Fig 4c). We found a weak enrichment for decreased turnover and increased stability in proteins whose gene deletions result in increased longevity (Fig 4c).

Extending our analysis to include all pro- and anti-longevity yeast genes reported in the GeneAge database^71^ revealed a subtle correlation between pro-longevity genes and decreased turnover and decreased stability (Extended Data Fig 3h). Importantly, longevity genes have been identified from single-gene deletion or single-gene overexpression studies, and there is currently no comprehensive understanding of genome-wide longevity traits. A recent genome-wide study of wild yeast isolates identified genes whose transcript abundance is positively (pro-longevity gene expression) or negatively (anti-longevity gene expression) correlated with RLS^72^. We found a strong association between pro-longevity gene expression and age-related increases in protein abundance and decreases in turnover, and the opposite pattern for anti-longevity gene expression (Fig 4d). Up-regulated proteins associated with pro-longevity gene expression were enriched in oxidative phosphorylation, overall energy production, and lipid metabolism, and down-regulated proteins associated with anti-longevity gene expression were enriched for glycolysis, amino acid, and nucleotide metabolism (Fig 4e).

Taken together, these findings suggest that age-dependent proteome remodeling from glycolysis/fermentation and amino acid metabolism to respiration and lipid metabolism is beneficial for aging and might contribute to longevity when modulated at basal levels. This is consistent with known pro-longevity interventions such as TOR pathway inhibition or caloric restriction, which lead to decreased amino acid biosynthesis or increased respiration, respectively^27^.

### Different molecular age-trajectories of protein complexes

Next, we asked whether our multidimensional proteomic atlas could elucidate age-dependent alterations in protein complexes, and enable a detailed understanding of how complex activity, assemblies, interactions, and biological functions change with age. As previously reported for different proteomic dimensions^21,24,73^, we find that subunits of the same annotated complex are highly correlated in their abundance, turnover, and stability compared to proteins that are not part of an annotated complex (Extended Data Fig 4a). Age scores revealed that larger protein complexes and proteins interacting with many other proteins had a tendency to decrease in abundance and stability and increase in turnover with age (Extended Data Fig 4b-c). This may suggest difficulties in maintaining the integrity of larger protein networks and protein complexes as a result of aging.

**Extended Data Figure 4.**
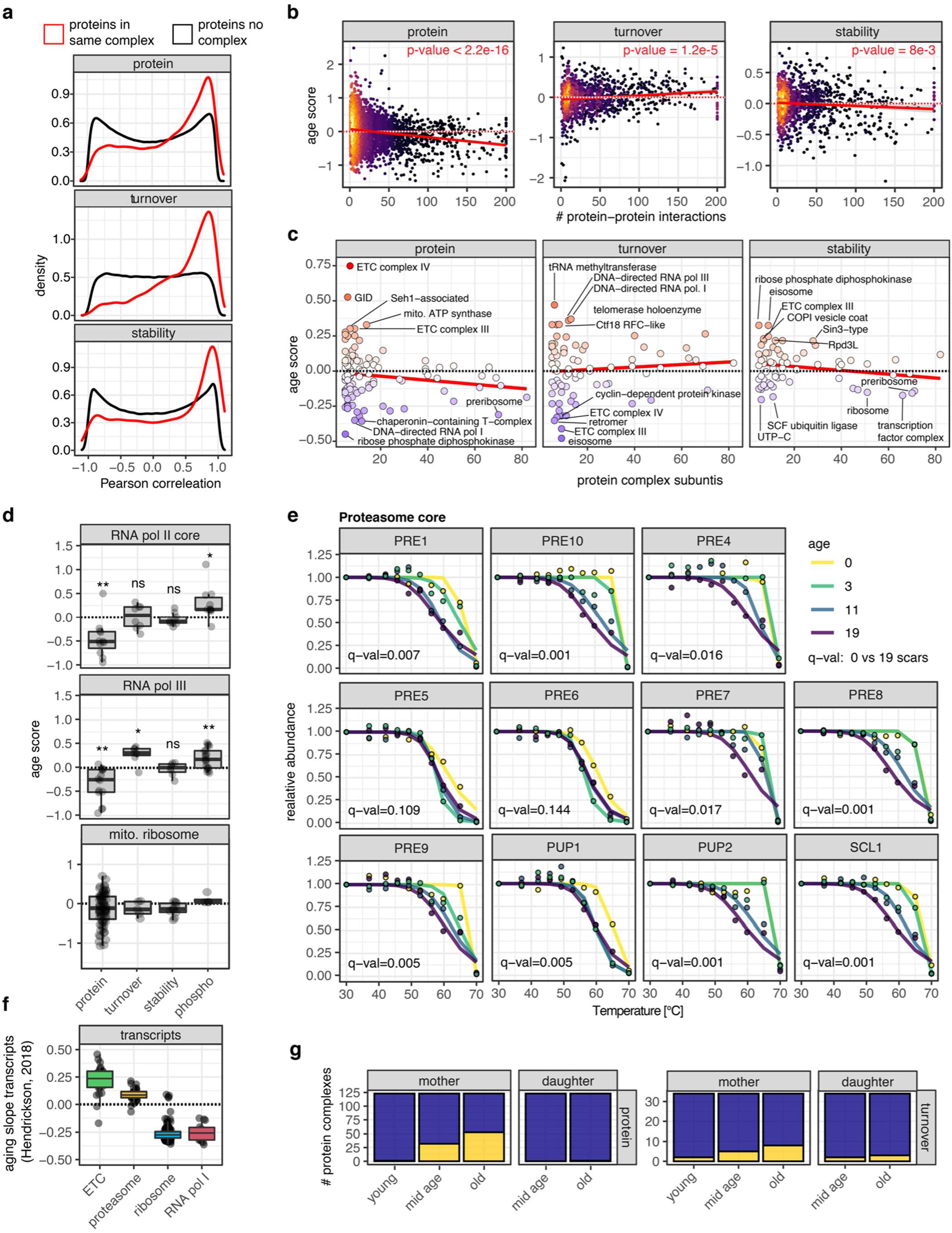
Age impact on protein complexes. **A)** Pearson correlation for proteins within the same complex and proteins not annotated to any complex for protein abundance, turnover and stability. **B)** Scatter plot of age score versus number of reported protein-protein interactions^95^ across different dimensions. **C)** Scatter plot of age score versus number of complex members across different dimensions. **D)** Age scores for subunits of the mitochondrial ribosome and for different subunits of RNA polymerase complexes. One-sided t-tests were performed to test whether group mean is different from 0 and statistical significance is indicated by: ns = non significant, * p< 0.05, ** p < 0.01. **E)** Melting curves for all measured subunits of the proteasome core complex. Q-value for young daughter versus old mother is indicated. **F)** Boxplots for aging slopes for transcripts of different protein complexes^49^. **G)** Bar plot of protein complexes, number of complexes that have more than 20% of subunits dysregulated are indicated.

We chose complexes of the electron transport chain (ETC), RNA polymerases, cytosolic ribosomes, and the proteasome for a more detailed analysis, since they showed strong but complex-specific age regulation (Fig 5a). The abundance of ETC complexes increased sharply, whereas turnover decreased and stability remained unaffected (Fig 5b). The abundance of DNA-directed RNA polymerase I subunits decreased, turnover increased, and multiple phosphosites were induced (Fig 5b, Extended Data Fig 4d). The abundance and stability of cytosolic ribosomal subunits decreased drastically, while their turnover increased (Fig 5b). In contrast, there was no discernible age-dependent trend for the mitochondrial ribosome (Fig 5d, Extended Data 5d). Multiple phosphosites on translation initiation factors eIF5 and eIF4B, as well as several ribosomal subunits showed an age-dependent increase (Fig 5b). Phosphorylation of these sites indicates translational inhibition and likely contributes to the known downregulation of protein synthesis in aged cells ^74^. The proteasome core and regulatory particle subunits showed heterogeneous protein abundance changes, decreased turnover, and the core subunits were severely and uniformly destabilized (Fig 5b, Extended Data Fig 4e).

**Figure 5.**
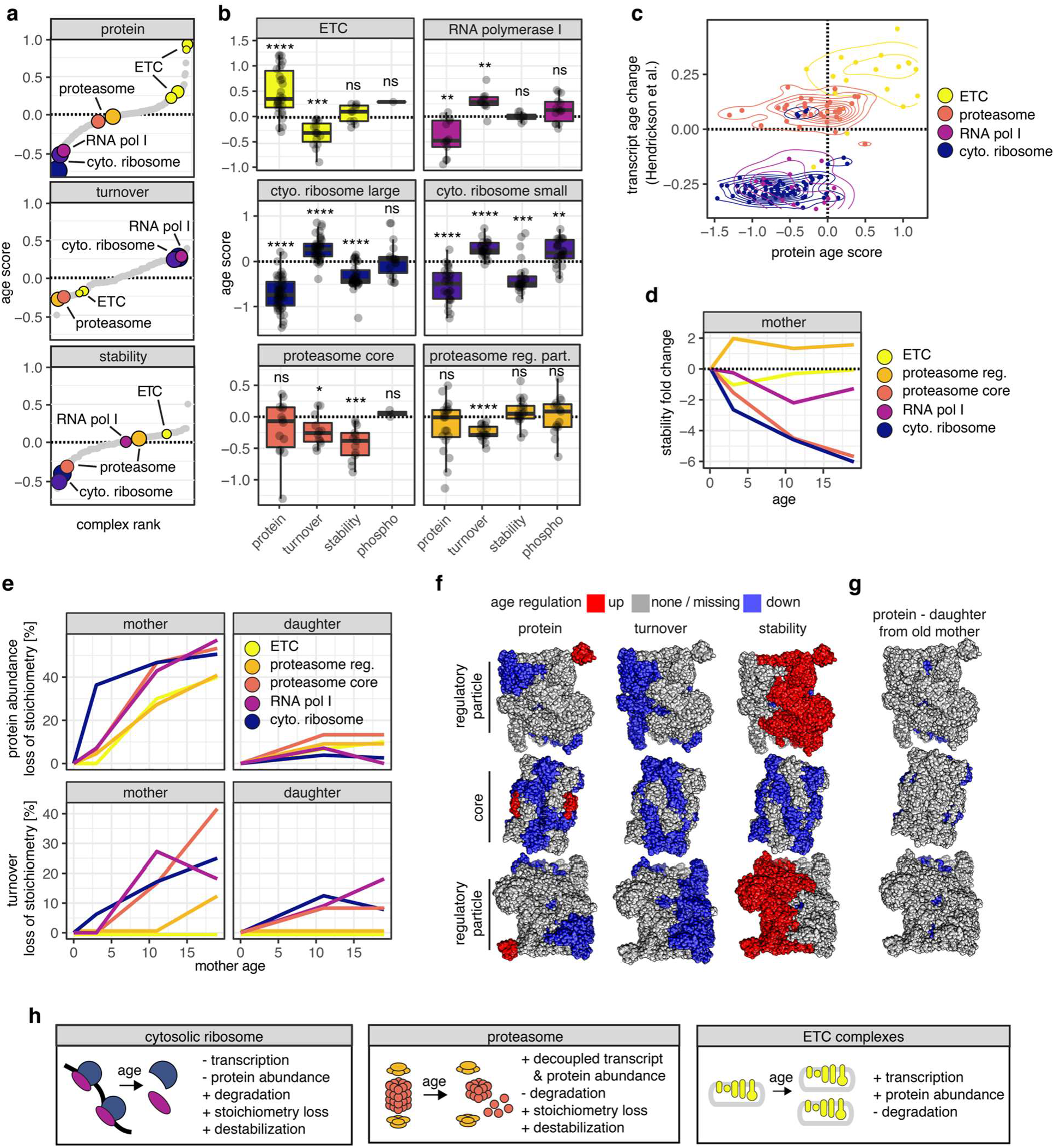
Dissecting age-trajectories of protein complexes. **A)** Age score rank plots for all annotated protein complexes. Selected complexes with extreme age-dependent behavior are indicated. **B)** Age score boxplot of all measured subunits for indicated complexes. One-sided t-tests were performed to test whether group mean is different from 0 and statistical significance is indicated by: ns = non significant, * p< 0.05, ** p < 0.01., *** p < 0.001, **** <p 0.0001. **C)** Contour plot for transcript age changes^49^ versus protein age score for selected complexes. Individual subunits are indicated by dots. **D)** Median fold changes of stabilities for selected complexes versus young daughter cells. **E)** Percentage of complex subunits that show stoichiometry loss for protein abundance (top) and turnover (bottom) for selected complexes. **F)** Mapping of significantly age regulated subunits onto the proteasome structure (PDB structures 4cr2 and 4qby). **G)** Same as F) for proteins regulated in daughters from old mothers versus young daughters. **H)** Model summarizing different protein complex age trajectories.

To determine whether specific complex regulation occurs post-transcriptionally, we compared age-dependent changes in subunit protein abundance with transcript abundance^49^ (Extended Data Fig 4f). We found good agreement for ETC, the ribosome, and RNA polymerase I, but strong disagreement for the proteasome (Fig 5c). Even though gene expression of proteasome subunits uniformly increased with age, changes in protein abundance were heterogeneous, and several subunits declined sharply, underscoring the importance of protein level readouts to account for post-transcriptional regulation.

Given the strong and differential regulation of subunit properties in major protein complexes, we evaluated if age-dependent changes in stability for selected protein complexes (Fig 5d) can be explained by potential changes in their stoichiometry. We therefore assessed complex stoichiometry dysregulation at the abundance and turnover levels^73^. The stability of the ETC and proteasome regulatory complexes was maintained, which was consistent with the observation that these complexes exhibited little to no dysregulation of subunit stoichiometry in the abundance and turnover dimensions, respectively (Fig 5e). The proteasome core and cytosolic ribosomes were severely destabilized, which could be attributed to their extensive stoichiometry dysregulation. We mapped proteasome subunits that were significantly affected by age onto the proteasome structure (Fig 5f). This confirmed strong age-dependent dysregulation of protein abundance and reduced turnover across the whole proteasome, and striking stabilization of the substrate interaction interface of the regulatory particle and global destabilization of the core. This suggests more proteasome substrate binding in older cells but no effective substrate processing due to a dysregulated core.

In general, the dysregulation of protein complex stoichiometry increased gradually with age, affecting around one-third of annotated protein complexes in old mother cells (Extended Data Fig 4g). Regardless of the age of the mother cell, protein complex stoichiometry was preserved or re-established in daughter cells (Fig 5e, Extended Data Fig 4g). Even for the highly dysregulated proteasome in old mothers, protein levels were re-established in their daughters (Fig 5g). The general decrease in proteasome turnover in old mother cells suggests the existence of mechanisms for the segregation of intact and newly translated proteasome complexes to daughter cells.

In conclusion, we observed various models of protein complex regulation in aging yeast that are controlled at the transcriptional, translational, and post-translational levels (Fig 5h). Both of the cytosolic ribosome subunits were affected by reduced gene expression, depletion at the protein level, and induced phosphorylation. The destabilization profile was indicative for a shift from 80s ribosomes and polysomes to 40s and 60s ribosomal subunits. This is consistent with previous studies that reported decreased translation of ribosome components, decrease in polysomes, and defects of translation initiation in old yeast cells^74,75^. Age-dependent decline of proteasomal function is a conserved feature in several species^76^ and age-dependent proteasome complex dysregulation has been reported before^50,77^. Our findings suggest a mechanism involving proteasome core destabilization and post-transcriptional stoichiometric loss. This is consistent with the pro-longevity effect of increasing overall proteasome levels in yeast^78^.

### Phosphorylation profiles and kinase activities across RLS

We identified more than 3300 phosphosites that were differentially regulated with age (Extended Data S5). We had previously classified the stress-responsive yeast phosphoproteome into functionally distinct signaling modules^25^. Applying this classification, we were able to associate age-dependent phosphorylation patterns with cellular stress responses and kinase activation states (Extended Data Fig 5a). Among the signaling modules that were most strongly regulated in aging, we highlight three: i) an age-downregulated TOR signaling module (21 phosphosites) that was associated with Tor (human MTOR), Sch9 (human SGK1), and PKA kinases; ii) an age-upregulated Snf1 signaling module (47 phosphosites); and iii) an age-upregulated MAPK signaling module (73 phosphosites) that was associated with MAPKs Ssk2 and Pbs2 belonging to the HOG pathway (Fig 6a, Extended Data Fig 5a). Comparing the age-dependent phosphorylation profile of these signaling modules versus stress-dependent profile showed that age induced signaling profiles were similar to those observed in carbon limitation, high pH, and high salt conditions (Fig 6b).

**Extended Data Figure 5.**
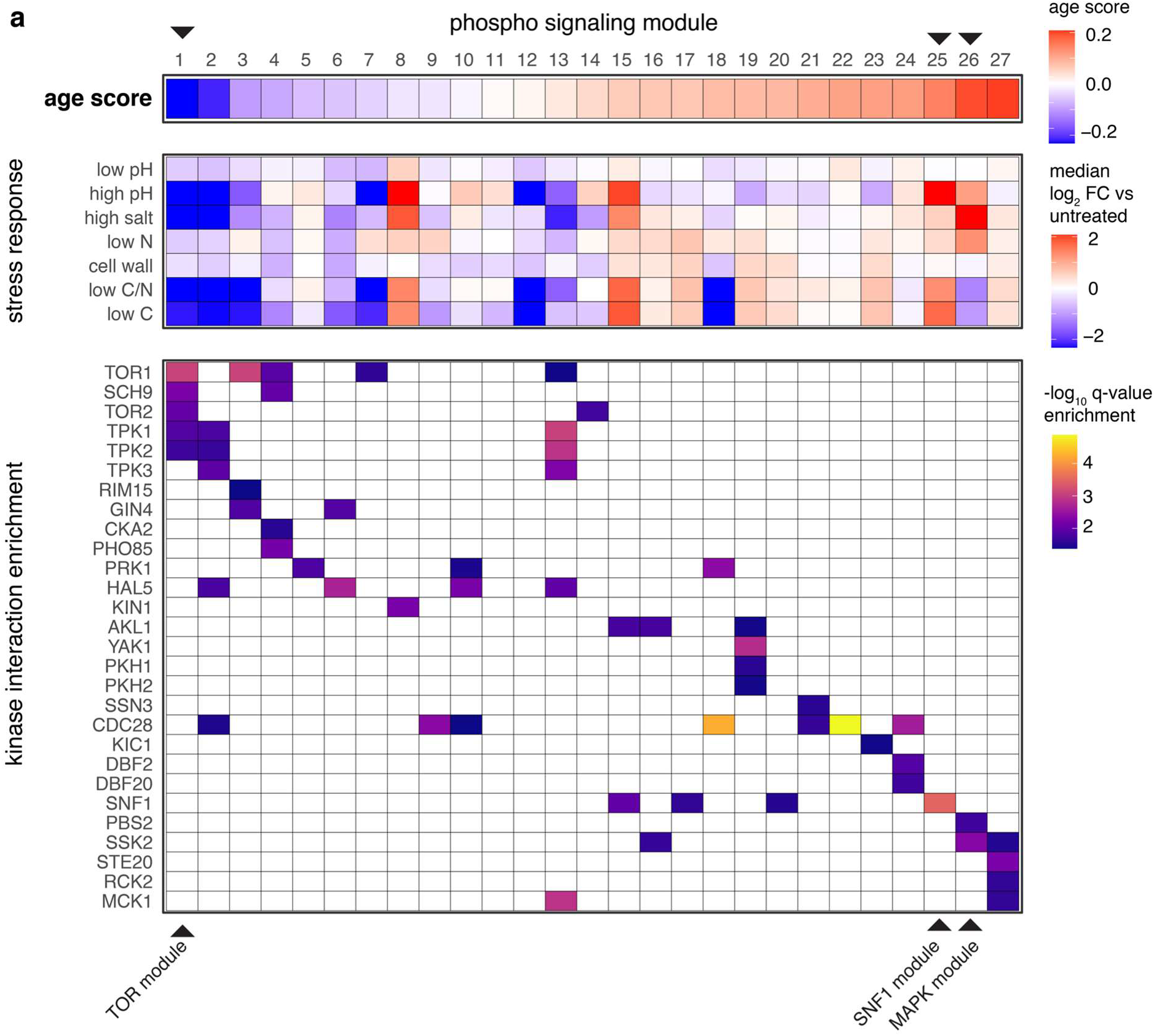
Age effect on phosphorylation signaling modules. **A)** 27 out of 29 phosphosite signaling modules^25^) that contain more than 10 phosphosites each are listed on the x-axis. Top panel: heatmap showing median phosphosite age scores for signaling modules. Signaling modules are ordered by increasing age scores. Middle panel: Heatmap of median fold change of phosphosites contained within signaling modules across different stress types^25^. Bottom panel: Heatmap of p-values for significantly enriched kinase-protein interactions within signaling modules. Black triangles on top indicate 3 signaling modules that are enriched for TOR, SNF1 or MAPK pathway components and were further analyzed in Fig 6a.

**Figure 6.**
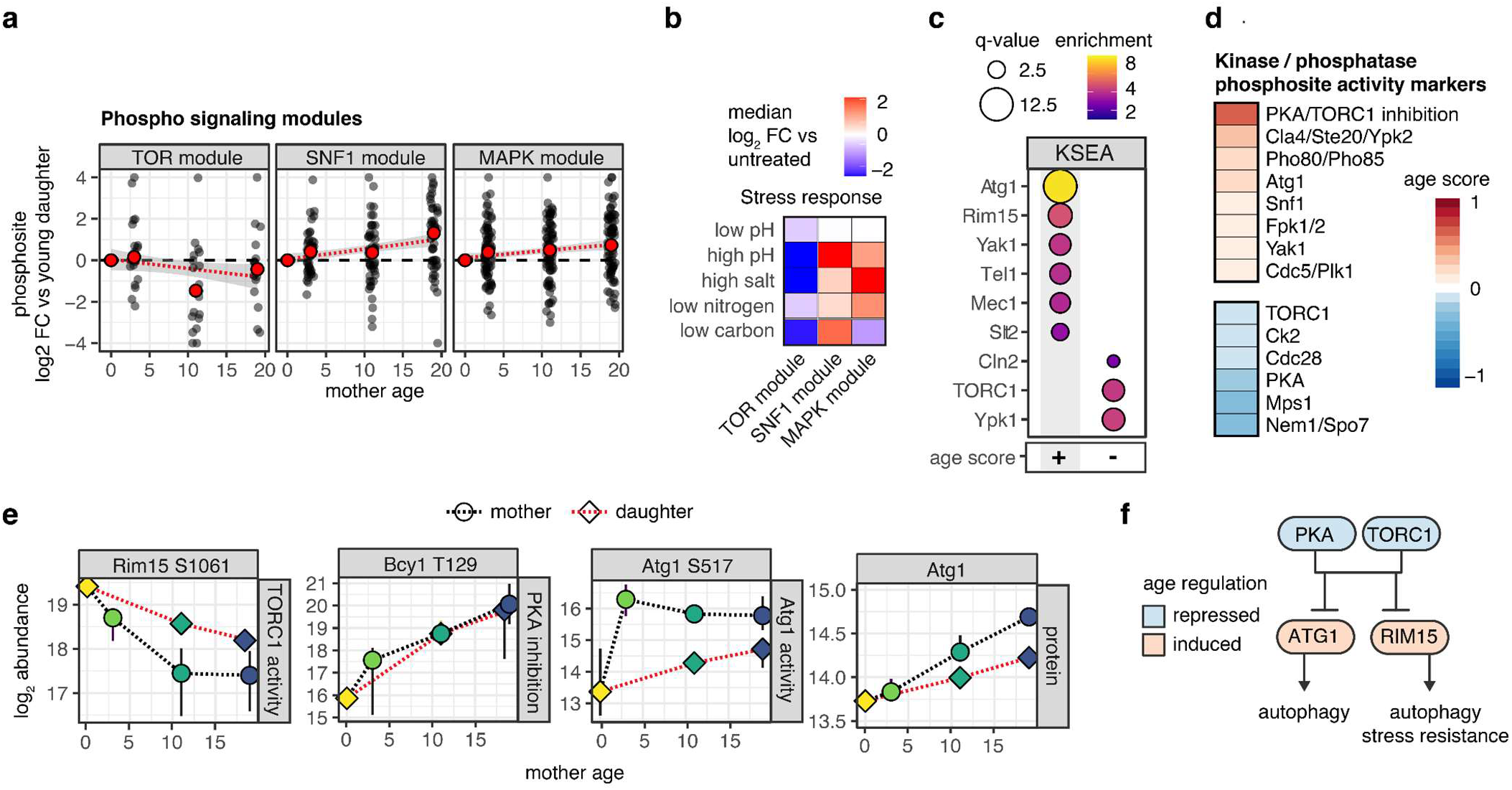
Age impact on kinase signaling pathways. **A)** Age-dependent regulation of phosphorylation sites belonging to different signaling modules^25^. Red dot indicates median phosphosite abundance and the red line corresponds to a linear fit. **B)** Behavior of selected signaling modules during stress responses^25^.**C)** Kinase-substrate enrichment analysis on phosphosite age scores. **D)** Median age scores for kinase activity phosphosite markers. **E)** Abundance of kinase activity phosphosite markers for Atg1, TORC1 and PKA. **F)** Model for age-effects on kinases involved in nutrient availability, autophagy and stress resistance signaling and their regulatory relationship.

To identify age-dependent activity changes of kinases, we performed kinase-substrate enrichment analysis (KSEA) using the age scores of known kinase target sites. We found age-dependent activation of autophagy kinases Atg1 (human ULK1) and Rim15 (human MASTL), genome integrity kinases Mec1 and Tel1 (human ATR and ATM), glucose sensing kinase Yak1 (human DYRK2) and cell wall integrity kinase Slt2 (human MAPK7) (Fig 6c). Kinases that showed age-dependent deactivation included TOR pathway kinases Tor1 and Ypk1 (human MTOR and SGK1) and cell cycle-promoting cyclin-kinase Cln2-Cdc28 (human CDK1) (Fig 6c). Furthermore, profiling age scores of well-defined kinase activity markers (e.g kinase activation loop phosphorylation)^79^ supported our KSEA results and additionally revealed an age-dependent inactivation of PKA and activation of Snf1 kinase (Fig 6d).

Next, we analyzed the behavior of key regulatory phosphosites within the TORC1 and PKA signaling network in order to understand how these signaling events change with age, and to interrogate the downstream consequences of these changes. Bcy1 is the negative regulatory subunit of PKA, and phosphorylation of Bcy1 on T129 results in its activation and inhibition of PKA^80^. Bcy1 is phosphorylated on T129 by Slt2 which in turn is inhibited through the TORC1-Sch9 axis^80^. We find gradual activation of Bcy1 as a function of age, both in daughter and mother cells, indicating broad PKA inhibition (Fig 6e). Rim15, a regulator of autophagy and cellular proliferation, is inactivated by phosphorylation on S1061 through the TORC1-Sch9 axis, which can be reversed by rapamycin treatment leading to increased RLS in yeast^81^ (Fig 6f). We revealed an age-dependent reduction of Rim15 S1061 phosphorylation in mother cells and to a lesser extent in daughter cells, consistent with TORC1 deactivation (Fig 6e). Atg1 autophosphorylation site S517 was asymmetrically induced in mother cells (Fig 6e). Additionally, Atg1 protein levels showed an asymmetric, age-dependent increase (Fig 6e).

Taken together, we find pervasive age-dependent changes in the phosphoproteome that suggest deactivation of TORC1 and PKA and activation of downstream autophagy and/or stress associated kinases Atg1 and Rim15 in aging (Fig 6f). Interventions that lead to inactivation of the TOR signaling axis and/or promote autophagy have been identified to increase lifespan in yeast and other organisms. However, the activity profile of these pathways in RLS is unclear. Strikingly, we found that TOR and autophagy signaling change in aging in a similar way as one would expect from pro-longevity treatments targeting these pathways. This finding supports a model that attributes the beneficial effects of TOR pathway inhibition to the rewiring of processes downstream of TOR in young cells, such as growth and autophagy, in anticipation of age-related deficiencies. It is possible that the reduction of TOR signaling in older cells is an effect of longer doubling time and eventual cessation of replication.

### Age-dependent and asymmetric activation of AMPK/Snf1 signaling and its consequences

To gain a systematic view of age-dependent rewiring of cellular pathways, we analyzed aggregated age scores for protein levels and phosphosite pathway activity markers^79^. Aging induced responses included glucose limitation, starvation, stress responses, and aerobic growth (Fig 7a, Extended Data Fig 6a). Aging-suppressed responses included anaerobic growth, amino acid starvation, and iron deficiency (Fig 7a, Extended Data Fig 6a). Age had a significant effect on the glucose sensing and signaling system, as measured both by protein abundance remodeling and phosphorylation rewiring (Fig 7a). As a consequence, the abundance of high-affinity hexose transporters increased with age, as did that of respiratory proteins, while the abundance of glucose gene repression mediators Mig1 and Hxk2 decreased (Fig 7b). These changes consisted of a clear shift from the high-glucose-fermentation to low-glucose-respiration gene expression program, which was also observed at the transcript level by Hendrickson et al.^49^. Protein changes were observed both in aging mother cells as well as their daughter cells, with the exception of the Snf1-Hxk2-Mig1 glucose sensing system, which was regulated in mother cells but reset in daughter cells. Snf1 (human AMPK) phosphorylates Mig1 at S311 and Hxk2 at S15, leading to their exclusion from the nucleus and released transcriptional repression from respiratory target genes^82^. Strikingly, we find strong asymmetry in mother versus daughter cell phosphorylation states of Mig1 and Hxk2, with high phosphorylation in mother cells, corresponding to derepression of respiratory genes (Fig 7c). These findings suggest that mother cells induce an age-dependent glucose limitation transcriptional program via activation of the AMPK/Snf1 energy sensor. The transcriptional program leads to increased abundance of proteins required for respiration and low glucose growth, which are inherited by daughter cells; however, AMPK/Snf1 sensing and signaling system is deactivated in daughter cells (Fig 7d).

**Figure 7.**
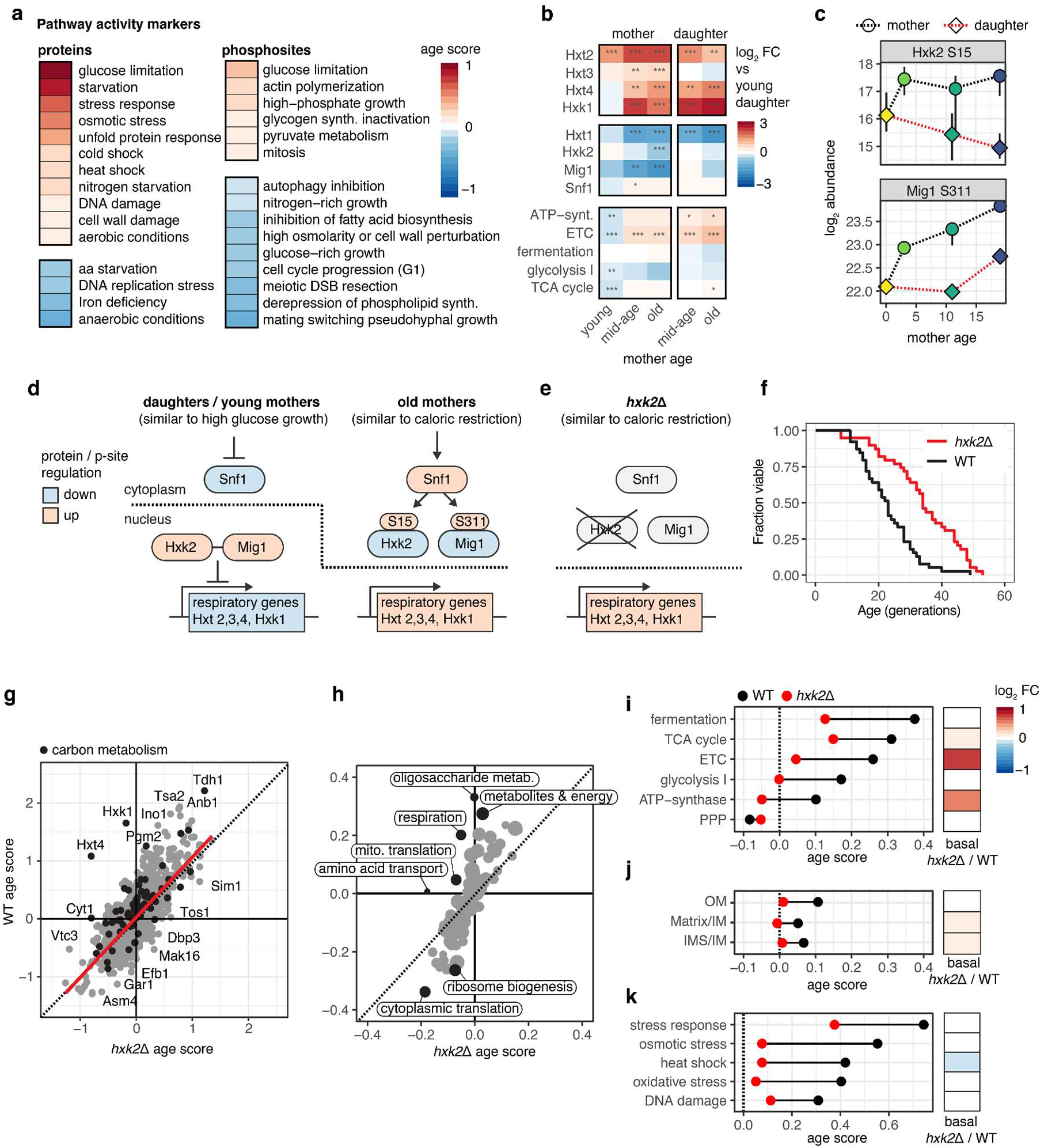
Reduced age-dependent proteome remodeling in a caloric restriction model. Median age scores for protein and phosphosite pathway activity markers. **B)** Protein abundance fold changes for regulatory and effector proteins involved in glucose repression. **C)** Abundance of regulatory phosphosites on Hxk2 and Mig1. **D)** Model of glucose repression system in young and old cells. **E)** Model of glucose repression system rewiring in *hxk2*Δ. **F)** Survival curves from microdissection experiments of WT and *hxk2*Δ strains (n= 40 cells per strain). **G)** Scatter plot of age scores for protein abundance in the WT vs *hxk2*Δ strain. The most different proteins are annotated and proteins involved in energy metabolism are marked in black. Red line corresponds to linear fit. **H)** Scatter plot of mean age scores for biological processes for WT vs *hxk2*Δ strain. The most different processes are marked in black and annotated. **I)** Left: Dotplot of median protein abundance age scores for proteins involved in energy metabolism for *hxk2*Δ in red and WT in black. Right: Heatmap depicting median log_2_ fold changes of proteins in *hxk2*Δ versus WT strains in young cells. **J)** Same as I) for proteins annotated with different mitochondrial sub localizations. **K)** Same as I) for proteins involved in different stress responses.

**Extended Data Figure 6.**
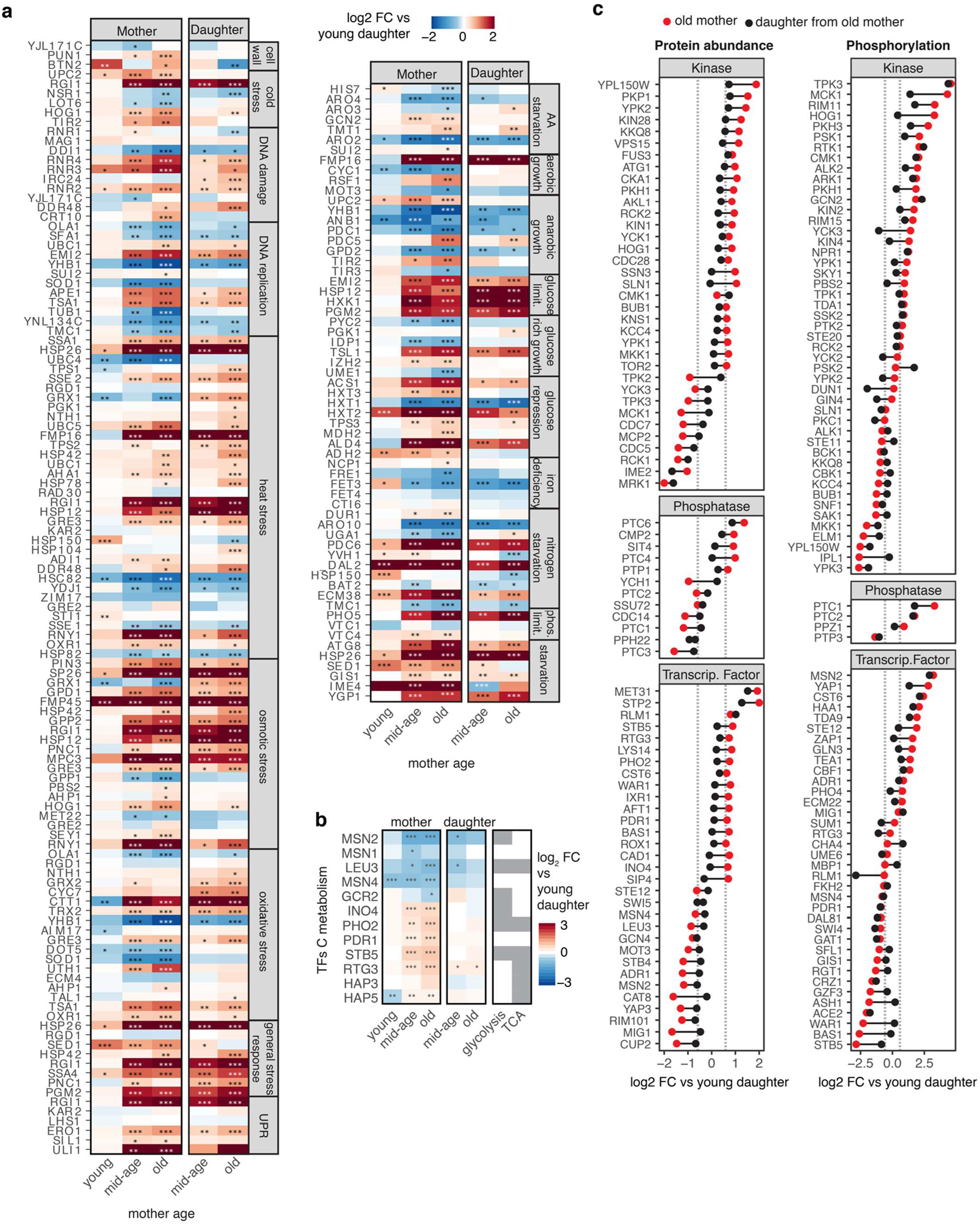
Age effect on stress markers, kinases, phosphatases and transcription factors. **A)** Fold changes versus young daughters for protein pathway markers. * q< 0.05, ** q < 0.01., *** q < 0.001, *** q < 0.001. **B)** Fold changes of carbon metabolism transcription factors versus young daughters. * q< 0.05, ** q < 0.01., *** q < 0.001, *** q < 0.001. **C)** Fold changes of old mothers and daughters from old mothers versus young daughters for protein abundance (left) or mean phosphosite abundance (right) for kinases, phosphatases and transcription factors that are significantly regulated in at least one condition.

Intriguingly, signaling reset in daughters appears to be a widespread mechanism, as we observe asymmetric mother-daughter regulation of protein abundance for virtually all carbon metabolism transcription factors (Extended Data Fig 6b). Prior studies suggested that aged yeast experience a failure in their ability to sense glucose properly, which is not due to an inability to import glucose but rather to an improper “information processing” of extracellular glucose levels^46,83^. Indeed, our analysis identifies differential activation of the AMPK/Snf1 energy sensing and signal transduction systems as the main age-dependent difference between mother and daughter cells. This also suggests that AMPK/Snf1 is induced by internal cues in mother cells, since mother and daughter cells are exposed to the same high-glucose environment.

We see asymmetric regulation of kinases, phosphatases and transcription factors involved in various other signaling pathways (Extended Data Fig 6c). For example, the master regulator kinase of osmotic stress, Hog1, is up-regulated in old mother cells and shows strong induction of phosphorylation in its activation loop, both of which are reversed in daughter cells (Extended Data Fig 6c). In summary, we identify widespread differences in signaling between mother and daughter cells.

### Reduced age-dependent proteome remodeling in a caloric restriction model

Numerous alterations that we found in aging, such as the relieving of glucose repression, the upregulation of respiratory metabolism, the inactivation of TOR and PKA signaling, and the downregulation of ribosome biogenesis, are similar to processes that are expected to take place during caloric restriction in yeast^84^. Caloric restriction is known to prolong the lifespan of many organisms, including yeast, by affecting various aging processes concurrently^85^. Most molecular studies of caloric restriction have been conducted on young cells, and the results correlated with the effects on lifespan in old cells. To directly test how caloric restriction affects the aging proteome and modulates the aging process, we turned to the *HXK2* deletion (*hxk2Δ*) model. By disrupting the entry of glucose into glycolysis and releasing glucose repression, *hxk2Δ* simulates the effects of low glucose and has been proven to likewise increase lifespan (Fig 7e)^86,87^. We validated the significant lifespan increase of *hxk2Δ^48^* using a RLS assay (Fig 7f). We compared the proteome of exponentially growing *hxk2Δ* and WT cells and confirmed our hypothesis that the *hxk2Δ* proteome showed similar alterations as we identified in old WT cells, with oxidative phosphorylation proteins, TCA cycle proteins and high-affinity glucose transporters being more abundant (Extended Data Fig 7a-b). Next, using WT and *hxk2Δ* strains, we conducted a MAD-proteomic time course similar to the previous experiments. Growth rates and median mother ages were comparable for WT and *hxk2Δ* (Extended Data Fig 7c). We performed quantitative proteomic measurements on young, middle-aged, and old mother and daughter cells from WT and *hxk2Δ* strains, determined differentially regulated proteins, and retrieved protein abundance age scores. We found a strong correlation between protein age scores in the two strains, with several proteins in the *hxk2Δ* strain being less affected by age (Fig 7g). Proteins involved in glycolysis, TCA, and oxidative phosphorylation showed strong age-dependent regulation in WT but much lower or no age-dependency in *hxk2Δ*. The comparison of average age scores for biological processes clearly showed that oligosaccharide metabolism and respiration were not affected by age in *hxk2Δ* (Fig 7h). Interestingly, *HXK2* deletion also led to an attenuated reduction of ribosomes and ribosome biogenesis machinery in aging (Fig 7h). By analyzing age effects on all energy-metabolic pathways in WT and *hxk2Δ* cells, we discovered that *hxk2Δ* cells had a significant reduction in age-dependent remodeling of all energy metabolic pathways, with proteins involved in oxidative phosphorylation and the TCA cycle being already high in young cells (Fig 7i). Moreover, the mitochondrial morphology of WT and *hxk2Δ* was comparable in young cells and remained unaffected by age in *hxk2Δ* (Fig 7j), which may delay age-dependent mitochondrial dysfunction. Interestingly, we also found a general dampening of multiple age-dependent stress responses in *hxk2Δ* (Fig 7k).

**Extended Data Figure 7.**
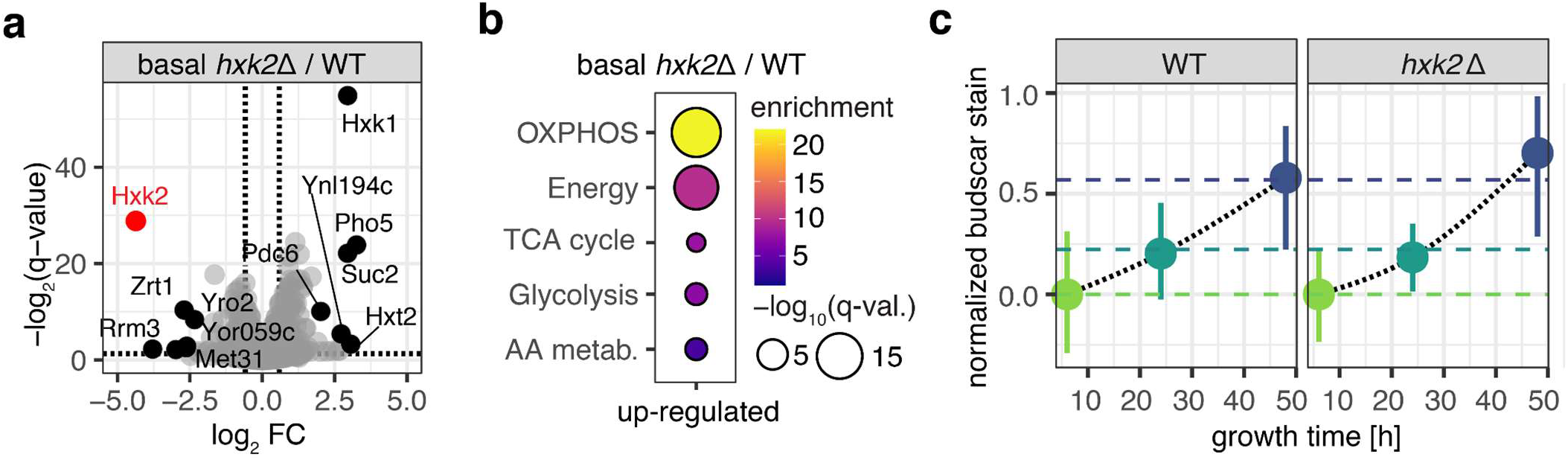
*Hxk2*Δ proteome and aging characteristics in ministats. **A)** Volcano plot of protein abundance in *hxk2*Δ vs WT under basal conditions (batch cultures, exponential growth). **B)** Proteome allocation enrichment for higher abundant proteins in *hxk2*Δ vs WT. **C)** Bud scar count of mother cells from *hxk2*Δ and WT strains derived from MADs.

In summary, we could differentiate between caloric restriction effects that influence the aging proteome in two major ways: i) young cells experiencing caloric restriction have already undergone some changes that occur during normal aging, this included rewiring of glucose metabolism to respiration, and; ii) changes that only affected older cells, which included improved maintenance of proteins involved in biogenesis, reduced morphological changes to the mitochondria, and reduced induction of stress responses. This suggests that the limited proteomic remodeling of energy pathways could contribute to conserving resources during aging and thus indirectly mitigate the decline in biogenesis, mitochondria, and response to stress.

## Discussion

We provide a multidimensional proteomic atlas of *Saccharomyces cerevisiae* across RLS, which can be explored interactively at https://rlsproteomics.gs.washington.edu. Our study revealed a substantial remodeling of the proteome with organismal age, reflecting alterations in metabolic activity, complex assemblies, post-translational regulation, signaling pathway activations, asymmetric cell division, and transcriptional programs. We demonstrate the utility of our integrated data analysis approaches for biological hypothesis generation and anticipate that it will become a valuable resource to study the basic biology of aging and conserved aging pathways.

We verified previously identified yeast aging hallmarks and characterized them further by defining the age of onset, proteome-wide impact across multiple dimensions, and transmission to daughter cells. Among these hallmarks, we mechanistically dissected mitochondrial dysfunction, a highly conserved aging pathway, at multiple levels that connected proteome alterations to morphology and function. In addition to a general increase in mitochondrial proteins, aged mother cells exhibited alterations in mitochondrial morphology. This was indicated by an increased ratio of mitochondrial surface to matrix proteins, along with age-dependent protein changes revealing an induced retrograde response, decreased mitochondrial fusion, and increased mitochondrial fission. Overall, these observations suggested mitochondrial fragmentation and a reduction of membrane potential. Concurrently, energy-metabolic pathways were remodeled toward respiratory metabolism. We identified the underlying regulation as an asymmetric activation of the AMPK/Snf1 energy sensor in mother cells, resulting in the relief of transcriptional glucose repression. While daughter cells from aged mothers continued to display altered mitochondrial proteomes, glucose repression signaling was reset at both the phosphorylation and transcription factor abundance levels in daughter cells. Since both mother and daughter cells are exposed to the same high-glucose environment, the asymmetric activation of the AMPK/Snf1 energy sensor in mother cells indicates that it responds to internal cues only present in the aged mother rather than actual extracellular glucose levels. One possible explanation for this observation is a previously proposed mechanism in which a gradual increase in yeast cell size at similar glucose influx rates leads to a decrease in intracellular glucose concentration ^50,51,88^. We also discovered that functional groups of the mitochondrial proteome were distributed asymmetrically between old mothers and their daughters, which is consistent with the hypothesis that daughters inherit healthier mitochondria (McFaline-Figueroa et al., 2011). Profiling of the mitochondrial proteome in a genetic caloric restriction model revealed that proteins of mitochondrial energy-metabolic pathways are already present in young cells at similar levels as in old WT cells and are not further regulated upon aging. This indicated an overall minimal remodeling of the mitochondrial proteome in the caloric restriction model, which is consistent with a recent study of yeast cells aged in low glucose conditions^89^. We further found that in the caloric restriction model, mitochondria were protected from age-dependent morphological changes. Together, these findings support a model in which mitochondrial dysfunction is associated, at least in part, with age-dependent remodeling of metabolic pathways and activation of AMPK/Snf1 signaling.

A prominent feature of yeast RLS is the asymmetric cell division that leads to the retention and accumulation of macromolecules in mother cells and the rejuvenation of daughter cells^30,31^. We found strong signatures of asymmetric protein segregation to mother cells as a result of aging in the protein abundance and protein turnover data, which show high overlap with previously reported asymmetrically partitioned proteins^33–36^. We identified 198 new asymmetrically segregated proteins by searching the entire proteome for proteins with increasing abundance and decreasing turnover signatures. Moreover, we found instances where asymmetric segregation broke down in old cells, which may contribute to the diminished rejuvenation of daughter cells derived from old mother cells ^40^. Failed asymmetric segregation predominantly affected proteins in the cytosol and mitochondria, and to a lesser extent proteins confined within organelle substructures known to distribute asymmetrically, such as the vacuole, the cell wall, and the plasma membrane. As mother cells grow significantly in size with age, it is likely that cytosolic diffusion barriers lose their effectiveness. This is supported by our observations of broad dysregulation of cytoskeletal proteins. Strikingly, our investigations revealed that asymmetric segregation also applies to protein complexes, protein phosphorylation, and signaling pathways. We find that the progressive dysregulation in complex stoichiometries and turnover in aged mother cells, which affected 30-40% of annotated complexes, is almost completely reversed in their daughter cells. The underlying mechanism could involve the retention of damaged, misfolded, aggregated, or incompletely assembled proteins or complexes in mothers; the restoration of protein homeostasis in daughters; or changes in subunit binding affinities due to cellular size increase and dilution. We anticipate that restoring the functionality of protein complexes is required for daughters to reach their full lifespan potential. Interestingly, we identify for the first time the asymmetric regulation of functional phosphosites implicated in multiple aging pathways, including in AMPK/Snf1 signaling as discussed above. We revealed that the plasma membrane H^+^-ATPase Pma1 asymmetrically occurred in its doubly phosphorylated, high activity form in mother cells but to a much lesser degree in their daughter cells. Mother-daughter asymmetry of intracellular pH has been identified as a crucial determinant of yeast aging by directly influencing mitochondrial functions^45,61^. In response to age-dependent increases in cell size and vacuole expansion, asymmetric phosphorylation of Pma1 may provide the cell with rapid means of pH regulation. Similarly, the master autophagy kinase Atg1 was asymmetrically autophosphorylated in mother cells, indicating a mother-specific induction of autophagy, possibly via TOR signaling inhibition. Additionally, we find asymmetric phosphorylation across stress signaling pathways, including mother specific activation of osmotic stress master kinase Hog1. Asymmetric regulation of signaling systems likely reflects age-dependent differences in the cellular environment, and as such offer an opportunity to modulate age-dependent cellular responses. Asymmetric cell division has also been observed in cells of higher eukaryotes, such as stem cells^90^. Asymmetric regulation of signaling systems and protein complexes might be involved in cell fate decisions and age-dependent deterioration of stem cell and regenerative functions.

The yeast RLS model has proven to be very powerful in understanding pathways that mediate longevity and are conserved between divergent eukaryotic species^57,91^. We found no clear correlation between age-dependent changes in any of the proteomic dimensions and pro-longevity genes identified through single-gene deletion or overexpression studies. This is not surprising since lifespan extension acts through different mechanisms, and a comprehensive understanding of genome-wide longevity traits is still lacking. Nonetheless, we identified two broad categories of age-dependent changes that are modulated by established longevity interventions. The first category consisted of instances in which the mechanisms of longevity interventions can be directly attributed to rescuing or stabilizing proteome functions that deteriorate with aging. Examples for this are the observed decline of proteasome stoichiometry and stability, which is consistent with the pro-longevity effects of genetically increasing expression of proteasome genes^78^; the observed decline in V-ATPase subunits in mother cells, which can be ameliorated by Vma1 overexpression and leads to increase lifespan ^45,62^; and our finding that proteins that gradually decreased in abundance during aging were strongly enriched for essential genes, which could explain a recent discovery that overexpression of essential genes disproportionately increased lifespan in yeast^92^. The second category consisted of instances in which age-dependent changes correlated with alterations already present in young cells under conditions or treatments that promote longevity. This included the observation of a strong positive correlation between age-dependent increases in protein abundance and pro-longevity gene expression identified in wild yeast strains^72^; remodeling of energy metabolic pathways towards respiration, as seen in caloric restriction^39,87^; deactivation of TOR signaling and consequential downstream effects, as observed in *tor1Δ*, *sch9Δ*, rapamycin treatment, and dietary restriction ^27,93^; and induction of autophagy through activation of Atg1, as observed in several lifespan prolonging interventions^94^. This category of proteomic changes likely represents the cellular response to age-dependent cues, and by inducing these changes already at a young age, cells will experience less proteomic remodeling during aging, which could conserve resources or reduce stress and make cells more resilient to age-related cues. Our study of age-dependent changes in a caloric restriction model confirmed remodeling of metabolic pathways in young cells, which then remained unchanged during aging. Only observed upon aging, we found that caloric restriction led to improvements in the maintenance of proteins involved in protein biogenesis, reduction in morphological changes to the mitochondria, and dampening of stress responses. These types of proteomic changes with interventions extending RLS could be considered a transformation from fitness-optimized proteomes, selected by rapid mitotic growth and meiotic capacity, to longevity-optimized proteomes. The pathways in this category are highly conserved and offer a variety of strategies for plausibly extending the lives of mammals, given that late-life fitness is not affected by selection.

## Methods

### Yeast strains

The strains used in this study were the S288C-derivative *Saccharomyces cerevisiae* haploid strain BY4741 (*MATa, his3Δ1, leu2Δ0, met15Δ0, ura3Δ0*) and BY4742 (M*ATα, hxk2Δ::KanMX, his3Δ1, leu2Δ0, met15Δ0, ura3Δ0*) from the Yeast Knockout Collection (YKO).

### Magnetic beading of cells

Yeast was grown overnight in synthetic medium complete, 2% glucose, 0.2%mannose (SMC+C). Overnight cells were diluted to an OD of 0.1 in SMC+C and grown until they reached OD 0.6. Cells were washed twice in phosphate buffered-saline pH 7.4 (PBS) and resuspended in PBS, 30% PEG 3350 at 5 OD/ml. 50 ODs of cells (∼7.5 x 10^8^ cells) are used as a starting amount to label cells for one MAD. Cells were sonicated for 15 seconds in a bench top water bath sonicator to break up any clumps and immediately mixed with activated magnetic beads.

To prepare activated magnetic beads 6.25mg of Sera-Mag SpeedBead hydrophilic carboxylate-modified paramagnetic particles (GE Life Sciences) were washed twice in MES buffer (500 mM 2-(N-morpholino)ethanesulfonic acid, pH 5). The beads are then resuspended in a solution containing MES buffer, 100mM N-hydroxysuccinimide, 35mM 1-ethyl-3-(-3-dimethylaminopropyl)carbodiimide hydrochloride and incubated at room temperature for 30 minutes. After incubation, bears were washed on a magnet three times with MES buffer and resuspended in PBS. Activated beads were immediately added to cells and incubated at room temperature for 30 minutes. After labeling, cells were washed three times in SMC+C on a magnet and resuspended to a density of 2 ODs/ml (∼3 x 10^7^) cells in SMC+C. All samples were checked by microscopy for efficient beading of 1-5 beads per cell and no cell clumping. Beaded cells were immediately loaded into MADs.

### Building, loading, running and harvesting MADs

The MADs were built as described in^49^. MADs were allowed to fill with SMC+C. The labeled yeast cells were added through the air inlet and the cells are given 5 minutes to attach to the magnets ringing the MADs before the pumps are turned on to a rate of approximately 1 replacement volume per hour. Typically 16 MADs were run in parallel. Every 12 hours and 1 hour before collection, the media and air pump were turned off and each MAD was taken out and vortexed, to remove any daughter cells which may have been stuck between the labeled cells. For protein turnover experiments, media in MADs and media reservoir were exchanged at once with SMC+C, heavy-labeled (^13^C6/^15^N2) lysine 2h before harvest. At each stated time point, whole MADs were harvested and labeled mother cells were further purified by washing 10 times on a magnet with SMC+C at room temperature. Non labeled cells were isolated from these washes and should represent daughter cells. For proteomic experiments following samples were derived from MADs: young mothers and their daughters (young daughters) after 6h growth; middle-aged mothers and their daughters after 24h growth and; old mothers and their daughters after 50h of growth. Samples were either snap frozen in liquid nitrogen for proteome analysis or placed in PBS, 1M Sorbitol, 4mM EDTA for bud scar counting, live/dead or organelle staining.

### Flow cytometry and microscopy of cells derived from MADs

Viability across the time course was estimated using propidium iodide and flow cytometry. Cells were diluted to 10^7^ cells/mL, stained with 1µL of 5mg/mL propidium iodide, incubated in the dark at room temperature for 30 minutes and washed twice with PBS before measurement. Age was qualitatively assessed using the bud scar stain Wheat Germ Agglutinin, Alexa Fluor 488 (WGA488). Cells from each point in the time course were fixed with 70% ethanol. The fixed cells were diluted to 10^7^ cells/mL and stained with 1µL of 5mg/mL WGA488 and washed twice with PBS before measurement. Age and viability were using the FL1 channel and FL3 channel of an Accuri BD C6 Csampler flow cytometer. For each measurement, more than 10,000 cells were counted.

For manual measurements of bud scars, cells were stained with WGA488 and imaged using a Leica DMI6000 inverted microscope. For each time point, the bud scars of more than 40 individual mother cells were manually counted from z-stack images.

### Replicative lifespan microdissection assay

Microdissection experiments to determine RLS were done as previously described^48^. Cells were patched onto SC + 2% Dextrose plates and allowed to grow overnight. Then, cells were arrayed, and virgin daughters were selected for use in the lifespan assay. New daughters were manually removed from mothers until mother cells die.

### Denaturing cell lysis, protein reduction, and alkylation

For denaturing cell lysis frozen cell pellets were resuspended in a lysis buffer composed of 8 M urea, 75 mM NaCl, and 50 mM HEPES pH 8. Cells were lysed by 4 cycles of bead beating (30 s beating, 1 min rest on ice) with zirconia/silica beads followed by clarification by centrifugation. Protein concentration was measured for every lysate by BCA assay and adjusted to 1 mg per ml. Proteins were reduced with 5 mM dithiothreitol (DTT) for 30 min at 55°C and alkylated with 15 mM iodoacetamide for 15 min at room temperature in the dark. The alkylation reaction was quenched by incubation with additional 10 mM DTT for 15 min at room temperature. Lysates were stored at −80°C until further processing.

### Whole proteome sample preparation

Denatured lysates containing 50 µg protein were diluted with 100 mM HEPES pH 8.2 to 1.5 M Urea and proteins were digested in-solution overnight at RT using endopeptidase Lys-C at 1:50 enzyme:substrate ratio. The next day digests were acidified to a final concentration of 1.5% TFA. Precipitation was removed by centrifugation, peptides were cleaned up by solid phase extraction on a µHLB Oasis 96-well plate (Waters), and eluates dried down by vacuum centrifugation.

25 µg of dried peptides were resuspended in 100 mM HEPES buffer pH 8.2, 30% acetonitrile (ACN), and were labeled with 125 µg of TMTpro 16plex isobaric label reagent (Thermo Fisher Scientific, Lot: V1310018) for 1h at RT. The reaction was quenched by addition of 5% hydroxylamine to a final concentration of 0.5% and 15 min incubation. The 30 samples were equally distributed across 2 sets of TMTpro plexes and for each plex the same control channel containing a pool of all samples was included. TMTpro channels corresponding to the same plex were pooled together and acidified to pH 3 with hydrochloric acid. Acidified peptides were desalted using Sep-Pak tC18 columns (Waters).

Offline pentafluorophenyl (PFP) reverse-phase chromatography was performed on each multiplexed sample using a XSelect HSS PFP 200 × 3.0 mm; 3.5 μm column (Waters) as described ^96^ using two buffers: (A) 0.1% trifluoroacetic acid (TFA), 3% ACN and (B) 0.1%TFA, 95% ACN. 48 fractions were collected and combined into 16 pooled fractions and lyophilized.

### Protein turnover sample preparation

Reduced and alkylated urea lysates were distributed in a 96-well plate (50 μg protein per sample), desalted and digested with Lys-C at 1:50 enzyme:substrate ratio using the R2-P1 (Rapid-Robotic Proteomics) protocol implemented on a KingFisher™ Flex (Thermo Fisher) magnetic particle processing robot as described before^56^.

### TPP sample preparation

Pelleted yeast cells were resuspended in native lysis buffer (50 mM HEPES pH 7.5, 75 mM NaCl, 2 mM MgCl2, protease inhibitors) and lysed by 4 cycles of bead beating (30 s beating, 1 min rest on ice) with zirconia/silica beads followed by clarification by centrifugation (10 min at 21,000 x g) to remove cell debris. Supernatant was collected, protein concentration was measured by BCA assay, adjusted to ∼2.5 mg/ml protein, and each sample was aliquoted into 10 PCR tubes on ice. PCR tubes were incubated on a thermal cycler in two phases: first, a 5 min incubation at 30°C for all tubes; second, a 5 min incubation spanning a gradient of 10 different temperatures per sample (30.0°C, 37.0°C, 42.1°C, 45.7°C, 49.2°C, 52.7°C, 56.2°C, 59.8°C, 65.0°C, 70.0°C) for 5 min. After temperature treatment, lysates were incubated at room temperature for 5 min. All samples were then centrifuged at 21,000 x g 4°C for 1 h. After centrifugation, 75 µl supernatant from each temperature-treated sample was taken and mixed 1:1 with denaturing buffer (9M urea, 10 mM DTT, 50 mM HEPES pH 8.9, 75 mM NaCl) and incubated at 55°C for 30 minutes. All samples were then incubated in the dark with 15 mM iodoacetamide for 30 min to alkylate cysteines and the reaction was quenched with 10 mM DTT for 30 min at RT. For each temperature, 50 µg of reduced and alkylated protein lysate was digested with LysC (1:50 enzyme:substrate ratio) overnight at RT.

Desalting, TMT labeling and offline fractionation was performed as described for whole proteome sample preparation with the following adjustments. 25 µg peptides were labeled with 100µg TMT11 plex (TMT10plex Lot VJ306784 and TMT11plex Lot VF298087). TMT channels corresponding to different temperatures of the same samples were pooled together and a control channel containing a pool of all 30°C treated samples was included. This resulted in 12 multiplexed samples which were individually fractionated using offline PFP reverse-phase chromatography. 24 fractions were collected and combined into 6 pooled fractions and lyophilized.

### Phosphoproteomic sample preparation

Reduced and alkylated urea lysates were distributed in a 96-well plate (200 μg protein per sample), desalted and digested with Trypsin at 1:100 enzyme:substrate ratio using the R2-P1 (Rapid-Robotic Proteomics) protocol implemented on a KingFisher™ Flex (Thermo Fisher) magnetic particle processing robot^56^. Phosphopeptides were enriched using the R2-P2 (Rapid-Robotic Phosphoproteomics) protocol using a Kingfisher Flex and Fe^3+^-NTA magnetic beads (PureCube Fe-NTA MagBeads, Cube Biotech)^56^.

### Mass spectrometry data acquisition

Peptides were dissolved in 4% formic acid (FA), 3% ACN and analyzed by nLC-MS/MS using an Orbitrap Eclipse Tribrid Mass Spectrometer (Thermo Fisher) or an Orbitrap Exploris 480 Mass Spectrometer (Thermo Fisher), both equipped with an Easy1200 nanoLC system (Thermo Fisher). Peptides were loaded onto a 100 μm ID × 3 cm precolumn packed with Reprosil C18 3 μm beads (Dr. Maisch GmbH), and separated by reverse-phase chromatography on a 100 μm ID × 35 cm analytical column packed with Reprosil C18 1.9 μm beads (Dr. Maisch GmbH), housed in a column heater set at 50°C, using two buffers: (A) 0.125% FA in water, and (B) 80% ACN, 0.125% FA in water at 400 nl/min flow rate.

For whole proteome and TPP sample analysis, TMTpro and TMT labeled peptides were separated by a 40-min effective gradient ranging from 11 to 35% ACN in 0.125% FA and analyzed online on a Eclipse Tribrid Mass Spectrometer. Data-dependent acquisition (DDA) using an MS3-based method with real time search and 2-second cycle time was used. First, MS1 data were collected using the Orbitrap (120,000 resolution; maximum injection time 50 ms; AGC 4e5; mass range 500-1800 m/z; charge states 2-6; dynamic exclusion 45 seconds). MS2 scans of the most intense precursors were performed in the ion trap with CID fragmentation (isolation window 0.7 Da; ion trap scan rate rapid; NCE 35%; maximum injection time 35 ms; AGC 5e4). Online real-time search^97^ was used to search spectra against the yeast FASTA database (digestion enzyme LysC; maximum 1 missed cleavage; static modifications TMT or TMTpro on N-terminus and lysines; Carbamidomethyl on cysteines; variable modification methionine oxidation; maximum 2 variable modifications). MS2 scans that passed custom RTS thresholds (Charge 2 precursors: Xcorr > 1; Charge 3+ precursors: Xcorr > 1.5; all charge states: PPM error < 25) were sent for MS3 quantification. The top 10 matching fragment ions were isolated using synchronous precursor selection^98^ followed by HCD fragmentation and collected for an MS3 scan in the Orbitrap (resolution 50,000; TurboTMT OFF; NCE 45% for TMTpro or NCE 55% for TMT; mass range 100-500 m/z; maximum injection time auto; isolation window 1.2 Da; AGC 1.5e5).

For turnover sample analysis, peptides were separated by a 60-min effective gradient ranging from 8 to 30% ACN in 0.125% FA and analyzed online on a Orbitrap Exploris 480 Mass Spectrometer. The BoxCarmax-data independent acquisition (DIA) method was used to measure intensities of light- and heavy-lysine labeled (^13^C6/^15^N2) peptides as described by Salovska et al.^99^. Briefly, the method allowed multiplexed acquisition at both MS1 and MS2 levels by integrating 22 m/z-wide BoxCar windows and 2.5 m/z-wide MSX scans covering the analytical m/z range of the corresponding BoxCar MS1 in combination with gas-phase separation strategy where each sample was injected 4 times.

For phosphoproteomic sample analysis, phosphopeptides samples were separated by a 60-min effective gradient ranging from 6 to 30% ACN in 0.125% FA and analyzed online on a Orbitrap Exploris 480 Mass Spectrometer. DIA-MS measurements were performed by acquiring 30 × 24-m/z (covering 438-1170 m/z) precursor isolation window MS/MS DIA spectra (30,000 resolution, AGC target 1e6, auto inject time, 27 NCE) using a staggered window pattern and optimized window placements as described^25^. Precursor spectra (60,000 resolution, standard AGC target, auto inject time) were interspersed every 30 MS/MS spectra.

### Mass spectrometry data processing, identification and quantification

For whole proteome and TPP analysis, DDA raw files were converted to mzML formats using msconvert, and MS/MS spectra were searched against *S. cerevisiae* S288C reference target/decoy protein sequence database using Comet (v2019.01.02) with recommended settings for linear ion trap mass analyzers^100^. LysC was selected as the digestive enzyme with a maximum of 2 missed cleavages, constant carbamidomethylation modification of cysteines (+57.0215 Da), constant TMT (+229.1629 Da) or TMTpro (+304.2071 Da) modification of lysines and peptide N-termini, and variable modifications of methionine oxidation (+15.9949 Da). Search results were filtered with Percolator (v3.01)^101^ to 1% false discovery rate at the precursor level. Precursor quantity was determined by extracting MS3 TMT reporter ion intensities using IsobaricQuant^102^. Precursors were filtered for matching identification with the real time search results. Isotopic TMT impurities were corrected using MSnbase^103^. MSstatsTMT^104^ with default parameters was used to consolidate precursors across different fractions; global peptide median normalization (only for whole proteome analysis); reference channel based normalization between different plexes (only for whole proteome analysis); and protein summarization using only non-redundant peptides. To generate protein melting curves and calculate a specific melting temperature (Tm) from the TPP data, relative abundance measurements per protein and temperature were scaled to the value measured at 30 °C and subjected to the global normalization procedure described in^20^.

For protein turnover analysis, spectral library generation and spectral library searches of DIA data, Spectronaut v.15 (Biognosys) was used as described by Salovska et al^99^. Precursor quantifications were exported from Spectronaut and filtered for a signal to noise ratio > 5 and > 4 data points per peak. Precursors were summarized to peptide quantifications by filtering for combined light and heavy precursor quantifications with maximal intensity across all samples. Relative protein turnover values were obtained by dividing heavy peptide intensity by light peptide intensity, taking the median turnover across all unique peptides belonging to a protein, filtering for at least two measured peptides per protein and performing quantile normalization. Relative protein turnover is therefore normalized to prolonged division time and overall reduced translation rates in old cells. The final dataset was filtered for proteins that were detected in at least two replicates per condition.

For phosphoproteomic analysis spectral library generation and spectral library searches of DIA data, Spectronaut v.15 (Biognosys) was used. A hybrid phosphopeptide spectral library was generated by combining the yeast phosphos reference spectral library generated in^25^ together with DIA data encompassing the quantitative phosphoproteomic measurements. Standard search parameters were used, including fixed modification of cysteine carbamidomethylation and variable modification of methionine oxidation and serine, threonine, and tyrosine phosphorylation. A PSM and peptide FDR cutoff of < 0.01 and a PTM localization site confidence score cutoff of > 0.75 were chosen. Spectral library searches were performed with the following adjustments to standard settings: decoy limit strategy was set to dynamic with a library size fraction of 0.1, but not less than 5000, a precursor FDR cutoff of < 0.01 was enforced by choosing the data filtering setting “Qvalue”, no imputation or cross run normalization was performed, a PTM localization site confidence score cutoff of > 0.75 was chosen, multiplicity was set to false, and PTM consolidation was done by summing. Quantitative phosphoproteomics analysis was performed at the phosphosite level across the data by summing peptide quantification values and median normalization across individual samples. Phosphosite quantifications were normalized to median abundance of the corresponding protein across a condition and phosphosites for which protein abundance was not quantified were discarded. A filter was applied to keep phosphosites that were present in at least 3 out of 5 replicates. Missing values were imputed with a shifted normal distribution on small values.

### Differential regulation analysis

For whole proteome differential regulation analysis, MSstatsTMT^104^ was used to test for significant changes in protein abundance across conditions (pairwise comparisons) using a linear mixed-effects model fit and moderated t-statistics. P-values were corrected globally using Benjamini-Hochberg correction. A protein was called as significantly regulated between two conditions if its absolute fold change was at least 1.5 and its adjusted p-value was less than 0.05.

To identify significantly altered protein thermal stabilities in a condition compared to the young daughter cells, we compared protein melting curves of the respective conditions versus protein melting curves in the young daughter cells using non-parametric analysis of response curves (NPARC)^105^. In order to estimate the null distribution for our dataset, we generated a permuted dataset of null F-statistics and used these null values to calculate empirical p-values for each protein melting curve comparison^106^. All p-values were corrected for multiple hypothesis testing using the Benjamini-Hochberg correction. Protein stability was called as significantly regulated between two conditions if its adjusted p-value was less than 0.05.

For protein turnover and phosphosite differential regulation analysis LIMMA (https://bioconductor.org/packages/limma/) was used on all samples at once with a model matrix including the age and cell type (mother or daughter) for each sample. Significant differential regulation was then calculated for each treatment against the young daughter samples, and p-values were corrected globally using Benjamini-Hochberg correction. A protein turnover value or phosphosite was called as significantly regulated in a condition if its absolute fold change between two conditions was at least 1.5 and its adjusted p-value was less than 0.05.

### Multiomic factor integration and age score calculations

The MEFISTO option from the MOFA framework was used as implemented in the MOFA2 R package (v.1.4.0)^67,107^. MEFISTO was performed on normalized quantifications of protein abundance, protein turnover, protein stability and phosphosites for young daughters, young mothers, middle-aged mothers and old mothers for WT cells or for normalized quantifications of protein abundance for young daughters, young mothers, middle-aged mothers and old mothers for WT and *hxk2Δ* strains. Different proteomic measurements or strains were defined as views and average bud scar counts for the different ages as temporal covariate. Standard options were used for model and training options apart from: the parameter “scale_views” was set to TRUE; “num_factors” was set to 4 and “convergence_mode” to slow. Both models trained on either the multidimensional WT aging dataset or the WT-*hxk2Δ* protein abundance dataset identified factor 1 to capture smooth variation along age. We extracted the weights from the model for underlying features across views for factor 1 and use them as age scores.

### Other bioinformatic analyses

If not specified otherwise, R version 4.1.0 (https://www.r-project.org/) with the “tidyverse” package collection (https://www.tidyverse.org) was used for all analyses. Heatmaps were done using the “pheatmap” R package (https://cran.r-project.org/web/packages/pheatmap/index.html).

Protein enrichment analyses were performed against the whole yeast proteome using a Fisher exact test. Kinase substrate enrichment analyses were performed against all identified and annotated phosphosites as a background using a Fisher exact test. The “run_enrichment” function of the MOFA2 package was used for enrichment analyses using age scores. For all enrichment analyses Benjamini– Hochberg multiple-hypothesis correction was applied and filtered for q-values < 0.05.

For all boxplots, the lower and the upper hinges of the boxes correspond to the 25% and 75% percentile, and the bar in the box to the median. The upper and lower whiskers extend from the largest and lowest values, respectively, but no further than 1.5 times the IQR from the hinge.

For all pointrange plots, the line corresponds to the 25% and 75% percentile, and the point to the median.

### Databases

The *S. cerevisiae* S288C reference protein fasta database containing the translations of all 6,713 systematically named ORFs, except “Dubious” ORFs and pseudogenes created on 05/11/2015 by SGD (https://www.yeastgenome.org/) was used for all searches. Gene Ontology terms, phenotype annotations and protein complex annotations were downloaded from SGD (https://www.yeastgenome.org/) on 04/05/2022.Yeast proteome allocation terms were obtained from^69,108,109^. Kinases and phosphatase phosphosite target annotation were downloaded from BioGrid (https://thebiogrid.org on 10/23/2020) and completed with data from^79,110–117^. Phosphoproteomic signaling modules were obtained from^25^. Pro- and anti-longevity genes were obtained from the gene age database (https://genomics.senescence.info/genes/index.html)71 (downloaded 10/20/2022).

## Acknowledgements

We thank Ian Smith, Matthew Berg, Julian Ramos, Alexander Hogrebe, Anthony Barente, Luana Paleologu, Noelle Fukuda, Alexis Chang, Sophie Moggridge, Bianca Ruiz, Devin Schweppe, and Miguel Martin-Perez for useful discussions and feedback.

## Author contributions

Conceptualization: ML, JV; Methodology: ML, JA, ARO, KH, MM, RAR; Investigation: ML, JV; Funding acquisition: ML, MK, JV, MD; Supervision: MK, MD, JV; Writing – original draft: ML, JV; Writing – review & editing: ML, JA, ARO, KH, RAR, MK, MD, JV

## Data and materials availability

The mass spectrometry proteomics data have been deposited to the ProteomeXchange Consortium via the PRIDE partner repository^118^.

Whole proteome data PXD039775 (Username, Password: 8I4WG2gh). Protein turnover: PXD039778 (Username, Password: W9Kss9t1). Thermal proteome profiling: PXD039823 (Username, Password: HfE9Fm7H). Phosphoproteomics: PXD039776, (Username, Password: Ty6oCTqM).

## Supplementary Materials

Supplementary Table 1: Protein quantifications across all samples.

Supplementary Table 2: Protein turnover across samples.

Supplementary Table 3: Protein thermal stability across samples.

Supplementary Table 4: Phosphosite quantifications across samples.

Supplementary Table 5: Differential regulation of protein properties.

Supplementary Table 6: Age scores for protein abundance, turnover, stability and phosphosites.

Supplementary Table 7: Proteins asymmetrically segregated to mother cells.

Supplementary Table 8: Protein quantifications across WT and *hxk2Δ* samples.

Supplementary Table 9: Age scores for protein abundance in WT and *hxk2Δ*.

